# An integrative single-cell atlas to explore the cellular and temporal specificity of neurological disorder genes during human brain development

**DOI:** 10.1101/2024.04.09.588220

**Authors:** Seoyeon Kim, Jihae Lee, In Gyeong Koh, Jungeun Ji, Hyun Jung Kim, Eunha Kim, Jihwan Park, Jong-Eun Park, Joon-Yong An

## Abstract

Single-cell technologies have enhanced comprehensive knowledge regarding the human brain by facilitating an extensive transcriptomic census across diverse brain regions. Nevertheless, understanding the cellular and temporal specificity of neurological disorders remains ambiguous due to the developmental variations. To address this gap, we illustrated the dynamics of disorder risk gene expressions under development by integrating multiple single-cell RNA sequencing datasets. We constructed a comprehensive single-cell atlas of developing human brains, encompassing 393,060 single cells across diverse developmental stages. Temporal analysis revealed the distinct expression patterns of disorder risk genes, including autism, highlighting their temporal regulation in different neuronal and glial lineages. We identified distinct neuronal lineages diverged across developmental stages, each exhibiting temporal-specific expression patterns of disorder genes. Lineages of non-neuronal cells determined by molecular profiles also showed temporal-specific expressions, indicating a link between cellular maturation and the risk of disorder. Furthermore, we explored the regulatory mechanisms involved in early brain development, revealing enriched patterns of fetal cell types for neuronal disorders, indicative of the prenatal stage’s influence on disease determination. Our findings facilitate unbiased comparisons of cell type-disorder associations and provide insight into dynamic alterations in risk genes during development, paving the way for a deeper understanding of neurological disorders.

## Introduction

The human nervous system undergoes a gradual and complex developmental process that spans several decades, beginning with embryogenesis and continuing through infancy, childhood, adolescence, and young adulthood. This prolonged developmental period involves the formation of a myriad of functionally distinct cell types, circuits, and regions. Recent advances in single-cell technology have surpassed conventional regional analyses to include comprehensive surveys of the entire human brain. Large-scale studies such as the Brain Initiative Cell Census Network have substantially aided in constructing detailed brain maps using single-nucleus RNA-sequencing analysis^1–4^. Despite such efforts, extant literature possesses limitations, including the lack of donor-specific diversity in terms of brain development and the absence of temporal dimensions in existing brain atlas studies. These gaps underscore the necessity for enhanced, comprehensive approaches to delineate the intricate developmental processes in the human brain.

The examination of cell types can elucidate the relevant pathology of neurological disorders, holding immense clinical significance. Since risk genes or genetic variants associated with neurological disorders are heterogeneous, understanding the cell type of converging their risk would comprehend our knowledge of the complex etiology and pathophysiological mechanisms underlying the disorder. De novo variants in autism disrupt neuronal genes that are overexpressed in prefrontal cortex during the mid-fetal period^5^, indicating that the risk perturbation at a molecular level precede its clinical onset. Furthermore, single-cell transcriptomics hold potential to suggest a novel molecular subtype not apparent through traditionally observed symptoms. A large-scale single-cell study of Alzheimer’s disease postmortem brains was able to identify a novel excitatory neuron cell type enriched in the cohesin complex and DNA damage response factors, and its association with cognitive impairment in the patients^6^. This single-cell transcriptome approach enunciates the fundamental characteristics of these disorders as well as facilitates the development of more effective and personalized treatments. Such findings underscore the imperative for comprehensive single-cell transcriptome studies to dissect the roles of neurological genes across diverse cell types, further enhancing our understanding of brain disorders This study aimed at presenting the brain transcriptome at the single-cell level (BTS), a single-cell atlas for developing human brains and evaluating the cellular and temporal specificities of neurological disorder genes. Data from eight previously published single-cell transcriptomics studies were integrated to curate a dataset comprising 114 human postmortem brain samples spanning the early fetal (7 gestational weeks) to late adulthood stages (90 years old). Our atlas offers an comprehensive resource for elucidating the developmental trajectory of the human brain and identifying various cell types that represent distinct temporal windows of development. Furthermore, leveraging this dataset, we conducted an in-depth exploration of cell type-specific genes implicated in neurological disorders and delineated the molecular mechanisms underlying neurodevelopmental disorders and age-related neurological conditions. This study advances human brain research and provides valuable insights into normal and pathological neurological processes throughout the lifespan.

## Materials and Methods

### Collection of single-cell transcriptome datasets

Single-cell and single-nucleus RNA-sequencing datasets were obtained in raw count matrices from publicly available sources. Our datasets include 114 samples from 80 donors obtained from the Allen Brain Map Cell Types Database, European Genome-phenome Archive (EGA), Gene Expression Omnibus (GEO), and Human Cell Atlas Data Portal (**Table S1**). Count matrices were collated from including Allen Brian Human_M1 10X data (https://portal.brain-map.org/atlases-and-data/rnaseq/human-m1-10x, n = 2), Braun et al.^7^ (EGAD00001006049, n = 19), Cameron et al.^8^ (EGAS00001006537, frontal cortex, and ganglionic eminence region, n = 3), Hardwick et al.^9^ (GSE178175, n = 2), Herring et al.^10^ (GSE168408, n = 24), Morabito et al.^11^ (GSE174367, n = 7), Nagy et al.^12^ (GSE144136, n = 17), and Zhu et al.^13^ (GSE202210, n = 6). For the Herring et al. dataset, cells annotated as poor-quality clusters in the original publication were excluded. In the Morabito et al. dataset, cells lacking sample information were excluded. For the Braun et al., dataset, sex information was inferred using XIST expression. Metadata for each sample was collected and harmonized in the format of Human Cell Atlas, including the source dataset, donor ID, sample ID, sequencing platform, library batch, sex, age, stage, race, hemisphere, brain region, postmortem interval (PMI), and diagnosis (**Table S1B**). Age information was formatted in days, with the fetus’s age described in terms of gestational age. Samples were categorized into 11 developmental stages based on the definition by Kang et al.^14^.

### Quality control and integration

The raw count matrices from distinct datasets were concatenated into a single dataset. Gene names were standardized using a gene symbol dictionary derived from NCBI and HGNC, excluding genes without valid annotations. Cells with a small count of genes (<50 genes) and a large proportion of mitochondrial genes (>30% of the total gene count) were filtered out. Doublet detection was performed using Scrublet^15^ (v0.2.1), and putative doublets were removed. To ensure sample balance across datasets, 50 samples from the Braun et al. dataset were randomly selected for analysis. Normalization and log transformation were conducted using Scanpy^16^ (v1.8.2). Highly variable genes are selected within each sample and merged to prevent batch-specific biases. This involves calculating highly variable genes within each sample, sorting them by the number of samples they are identified in, and selecting the top 5,000 genes found in the most samples. Datasets were merged across different samples using scvi-tools^17^ (v1.0.3) and adjusted the batch effect by setting the batch key as the sample ID and additional categorical covariate keys including the source dataset, assay information (single-cell or single-nuclei), and library kit (10x 3’ v2 or 10x 3’ v3). The latent representation for each cell was used to compute the nearest neighbor distance matrix and construct a neighborhood graph. The Leiden algorithm was used for clustering with a resolution of 0.6. The weights from the scVI model were converted to a shareable trained model for further use with the single-cell architectural surgery (scArches) algorithm^18^.

### Annotation of clusters

The major cell types were identified based on annotations from original studies and expression profiles of cell type marker genes defined by the Allen Brain Institute^19^: SLC17A7 for excitatory neurons; GAD1 for inhibitory fetal; FGFR3 for astrocytes; TYROBP for microglia; OPALIN for oligodendrocytes; PDGFRA for OPC; NOSTRIN for endothelial cells; HES1 and SOX2 for radial glia; and NHLH1 and NEUROD6 for neuroblasts. Cell subtypes for neurons were defined by layer markers for excitatory neurons (LINC00507 for layers 2-3; RORB for layers 3-5; FEZF2, and THEMIS for layers 4-6) and branch markers for inhibitory neurons (PVALB and SST for Medial Ganglionic Eminence (MGE); LAMP5, VIP and ADARB2 for Caudal Ganglionic Eminence, (CGE)). Each cluster was further annotated by determining the most cluster-specific marker, exhibiting the greatest fold change with a false discovery rate (FDR) <0.05 compared to all other cells detected in at least 25% of the cells within the cluster (**Table S2**). The Wilcoxon rank-sum test was used for differential testing.

### Gene set enrichment test for cell type validation

A gene set enrichment test was performed for the established cell type marker genes from the previous single-cell transcriptomic study of the developing brain^20^. The background genes for the enrichment test were set to include all the genes included in the single-cell dataset of the previous study. Marker genes for each annotated cluster EN (excitatory neurons from postnatal samples), EN-fetal-early (excitatory neurons from early fetal samples), EN-fetal-late (excitatory neurons for late fetal samples), CGE-derived inhibitory neurons (IN-CGE), MGE-derived inhibitory neurons (IN-MGE), astrocytes, microglia, oligodendrocytes, OPC, pericytes, VSMC, endothelial cells, RG, and IPC were collected from the supplementary data provided. This enrichment test included cluster-specific differentially expressed genes (DEGs) with a threshold of FDR <0.05, log_2_ fold change >0.2, and >25% of cells within the cluster expressing the gene. A one-sided Fisher’s exact test with multiple comparisons was applied.

### Gene set enrichment test for neurological disorders and glioblastoma

A comprehensive set of risk genes associated with various neurological disorders and conditions identified in large-scale exome studies or genome-wide association studies (GWAS) was systematically collated. The risk genes were subjected to enrichment tests with cluster-specific DEGs with a threshold of FDR <0.05 and log_2_ fold change >0.2. A one-sided Fisher’s exact test with multiple comparisons was applied.

Risk genes of neurodevelopmental disorders, including autism, epilepsy, or developmental delay were selected. For autism, we utilized risk genes identified in large-scale exome studies. 185 genes with the enrichment of protein-truncating variants (PTVs), missense variants, and copy number variants (CNVs) in 20,627 autism cases qualified multiple comparisons at FDR <0.05 were selected from Fu et al.^21^. In addition, we utilized 373 risk genes from the meta-analysis of cohorts ascertained for developmental delay (DD). We also chose 102 genes that are enriched for 11,986 autism cases at FDR <0.1 from Satterstrom et al.^22^. Epilepsy risk genes were sourced from the Epi25 dataset, comprising a compilation of 20,979 cases and 33,444 controls from 59 global research cohorts. A total of 140 genes were selected from the summary statistics (https://epi25.broadinstitute.org/results) based on the enrichment of PTVs or damaging missense variants in case subjects (p-value <0.01).

For neuropsychiatric disorders, we chose the risk genes for anxiety disorder, bipolar disorder, major depression, and schizophrenia. We selected 692 genes from the anxiety-associated genomic loci in the GWAS catalog (EFO ID: EFO_0006788). We selected 58 bipolar disorder risk genes from the BipEx dataset, which encompasses data from 14,210 cases and 14,422 controls. Genes significantly enriched for PTVs or damaging missense variants (p-value <0.01) in cases were subset from the summary statistics (https://bipex.broadinstitute.org). We obtained 450 major depression-associated genes from a GWAS analysis of 88,316 cases and 902,757 controls^23^. Schizophrenia-related genes were sourced from exome and GWAS analysis. We utilized the schizophrenia exome meta-analysis consortium (SCHEMA) dataset, which includes data from 24,248 cases, 97,322 controls, and 3,402 parent-proband trios. Gene selection (n = 32) was guided by FDR <0.05. Additionally, 287 schizophrenia-associated loci and 2,132 genes were obtained from Trubetskoy et al.^24^, involving 76,755 individuals with schizophrenia and 243,649 controls.

For neurodegenerative disorders, we focused on genes associated with Alzheimer’s and Parkinson’s diseases. We selected 76 genes associated with Alzheimer’s disease from a recent GWAS comprising 111,326 cases and 677,663 controls^25^. Parkinson’s disease genes (n=423) were retrieved based on GWAS loci reported in the GWAS catalog (EFO ID: MONDO_0005180).

Furthermore, genes associated with neurological conditions in response to trauma exposure (EFO ID: EFO_0008483), vascular brain injury (EFO ID: EFO_0006791), and abnormal brain morphology (EFO ID: HP_0012443) were included. These were identified from GWAS loci reported in the GWAS catalog, with a specific number of undisclosed genes.

For glioblastoma, 17 glioblastoma driver genes were selected^26^. Gene set signatures for specific cellular states in glioblastoma (Astrocyte-like, OPC-like, NPC-like subprogram 1, NPC-like subprogram 2, Mesenchymal-like hypoxia-independent, and Mesenchymal-like hypoxia-dependent) were obtained from the meta-module gene list, identified to be recurrent across tumors indicating global characterization of intra-tumoral heterogeneity. Gene set signatures related to the cell cycle were also obtained for phases G1/S and G2/M for cell cycle-specific characteristics maintained across different tumor types^27^.

### Pseudo-time and trajectory analysis

Pseudo-time analysis was performed using Palantir^28^. Subset of each cell type of interest was reprocessed before the analysis. Samples with <100 cells were excluded from the pre-processing and integration steps to ensure robustness. To mitigate the batch effect, 5,000 highly variable genes were selected from each sample and integrated using scvi-tools^17^.

Diffusion maps were derived from batch-corrected embeddings, and the resulting components were projected onto Uniform Manifold Approximation and Projection (UMAP). Pseudo-time computation and trajectory construction were conducted by designating cells with the minimum age of the premature cell type as the initial cell for each group. For instance, in the neuronal group, one of the earliest radial glial cells was chosen as the initial state. Similarly, in the oligodendrocyte and astrocyte groups, one of the OPC cells and astrocytes with the minimum age was considered as the initial states. The endpoints of the trajectories were determined automatically. Differentiation potential was estimated by quantifying the entropy and pseudo-time distance of each cell from the initial state. Gene trends were subsequently computed for each lineage, and gene expression was illustrated across pseudo-time for temporal investigation.

### Analysis of gene expression profiles across developmental age

Pseudo-bulk aggregation of the expression matrix was performed using decoupler^29^ (v1.5.0). Expression profiles were summarized across cells per sample ID and Leiden cluster by calculating the mean log-normalized count. To ensure data quality, samples harboring a minimum of 10 cells and 1,000 accumulated counts were exclusively considered for the pseudo-bulk aggregation. Trend lines for sample-wise expression were generated by fitting using the Loess function with a span of 0.4 and enabling a smooth representation of expression trends across developmental ages. Leiden clusters containing more than 4,600 cells were included to capture robust population dynamics, and cells from the Nagy et al. dataset were excluded due to the unknown exact age of the samples.

### Inference of gene regulatory networks and enriched signaling pathways

To facilitate gene regulatory network inference, transcriptionally similar cells were categorized into meta-cells using SEACells^30^ (v0.3.3). Following the designation of one meta-cell for every 75 single cells, 393,060 cells were aggregated into 5,000 meta-cells. In the initialization step, the embedding matrix generated by scVI was used to compute the kernel for the meta-cells and prioritize the top 10 eigenvalues. Subsequently, we constructed the kernel matrix and conducted an archetypal analysis. In the model fitting step, the minimum and maximum iterations were set to 10 and 100, with a convergence threshold of 0.01125. Within each SEACell, cellular aggregation was achieved by summing the log-normalized expression of all constituent cells, generating aggregated counts.

Transcription factor regulatory networks were inferred using pySCENIC^31^ (v0.12.1). Log-transformed counts from the SEACells were used as an input matrix, considering only protein-coding genes. Adjacencies between the transcription factors and their targets were inferred using the GRNBoost2 algorithm. Regulon prediction for 1,892 transcription factors was performed based on the motifs of the transcription factors and putative promoter regions of the target genes obtained from the cisTarget database (version 9), covering 10kb around the transcription start site, 500 bases upstream, and 100 bases downstream. The correlation between transcription factor and target genes was calculated using the entire set of cells, including those with zero expression. Cellular enrichments of regulons per SEACell were determined with an AUC threshold of 0.05, and the resulting regulon activities calculated for each SEACell were matched to the Leiden clusters possessing the highest cell counts. The regulon activities for transcription factors with a minimum mean of 0.02 and a variance of 0.001 across Leiden clusters were visualized as a heatmap using the R package ComplexHeatmap (v2.15.1). Gene sets representing hormonal regulation, kinase-mediated pathways, and immune signaling pathways were obtained from the Reactome database^32^. Module scores were computed by averaging the expression levels of genes within each gene set and subtracting the average expression of a reference set of genes.

## Results

### Generation of a single-cell atlas for developing human brains

We established a single-cell atlas for developing human brains by compiling single-cell RNA sequencing (scRNA) and single-nucleus RNA sequencing (snRNA) datasets from eight studies comprising 80 donors and 114 postmortem brain samples (**Figures 1A, 1B, S1**). This comprehensive single-cell atlas integrates expert-curated, quality-assured, and pre-analyzed datasets from publicly available studies on the pre- and postnatal periods of the human brain. We selected samples that had not been previously reported for neurological disorders, assuming neurotypical brains. The samples encompassed a wide range of developmental stages from 7 gestational weeks to 90 years of age, and diverse brain regions (**Figures 1C, 1D**). As the datasets were heterogeneous in terms of data quality, gene name representation, and sample metadata, we rigorously conducted a quality control process for the datasets and created a consensus notation for the sample metadata (**Table S1**). To mitigate the batch effect when combining these datasets, we matched gene names across the datasets and integrated the raw matrix files of the single-cell datasets using the scVI tool^17^. Overall, the single-cell atlas integrated 393,060 single cells and 41 clusters, which were annotated as 10 major cell types and 22 cell subtypes, based on the Allen Brain Institute^19^ (**Figures 1E, 1F, 1G**).

**Figure 1.**
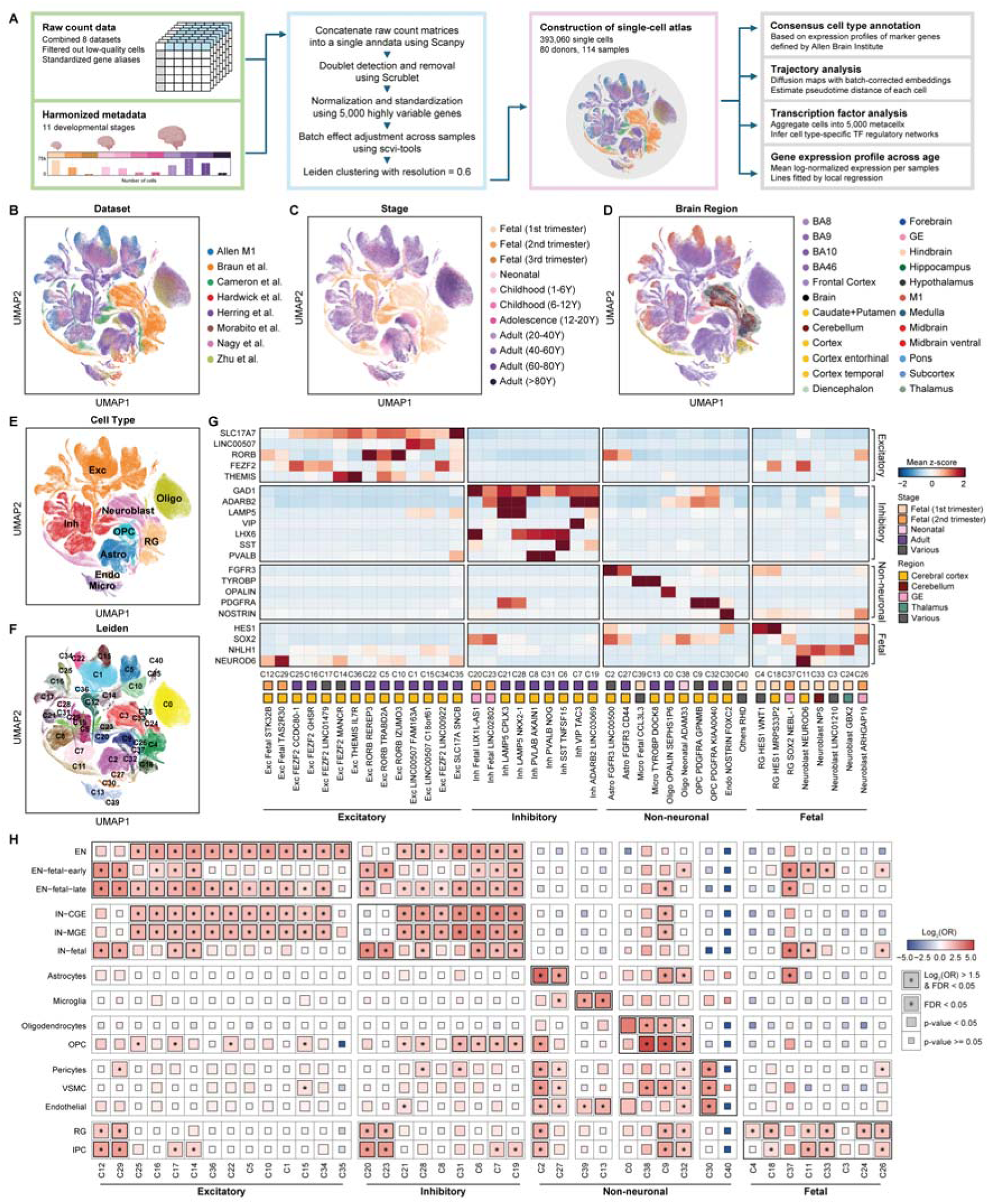
Integrated single-cell atlas of developing human brain. **A.** Schematic of atlas construction and downstream analysis. **B-F.** Uniform Manifold Approximation and Projection (UMAP) of the atlas, colored by (B) integrated datasets, (C) developmental stages, (D) brain regions described in original studies, (E) major cell types, and (F) Leiden clusters. **G.** Taxonomy of 41 Leiden clusters based on the scaled expression of marker genes. The stage and region with the highest proportion of cells (>35%) were designated. **H.** Gene set enrichment test for the established cell type marker genes sourced from a previous single-cell transcriptome study^20^. A one-sided Fisher’s exact test was used to compute statistics with multiple comparisons by FDR.

Regarding excitatory neurons, we identified cell subtypes based on developmental stages and cortical layers. We characterized 11 clusters of cortical layer-specific excitatory neurons expressing markers for the cortical layer 2-3 (LINC00507), layer 3-5 (RORB), and layer 4-6 (FEZF2 and THEMIS). Consequently, there are two clusters (C1 and C15) for layer 2-3, three clusters (C5, C10, and C22) for layer 3-5, and six clusters (C14, C16, C17, C25, C34, and C36) for layer 4-6. Additionally, we identified two clusters that are predominantly present in the fetal second trimester (C12 and C29). C12 displays a dynamic composition spanning from the 14 post-conception weeks to 35 months after birth, which could offer insights into the fetal-to-neonatal transition of excitatory neurons. Inhibitory neurons were classified into cell subtypes based on marker expression of the caudal ganglionic eminence (CGE) branch (ADARB2, LAMP5, and VIP) and medial ganglionic eminence (MGE) branch (LHX6, PVALB, and SST). We characterized seven clusters of branch-specific inhibitory neurons. In the CGE branch, two clusters (C21 and C28) expressed LAMP5, one cluster (C7) expressed VIP, and one cluster (C19) expressed ADARB2. In the MGE branch, there were two clusters (C8 and C31) expressing PVALB and one cluster (C6) expressing SST. Furthermore, we identified two clusters enriched in fetal inhibitory neurons (C20 and C23) that were prevalent in the second trimester. These clusters exhibited distinct expression patterns of branch markers. C20 was distinguished by the expression of LHX6, whereas C23 exhibited the expression of ADARB2, indicating their unique developmental origins.

Our single-cell atlas delineates clusters specific to various developmental stages. Clusters of radial glia (C4, C18, and C37) and neuroblasts (C3, C11, C24, C26, and C33) were predominantly present during the fetal first and second trimesters. Microglia and oligodendrocytes were subdivided into two clusters, each based on the developmental stage. We found that one microglial cluster was prevalent in the adult stage (C13), and the other was predominant in the fetal first trimester (C39). For oligodendrocytes, one cluster prevailed in the adult stage (C0), whereas the other predominated in the neonatal stage (C38). Astrocytes and OPCs were subdivided into two clusters. One cluster (C2 for astrocytes and C9 for OPCs) exhibited a mix of developmental stages, whereas the other cluster (C27 for astrocytes and C32 for OPCs) predominantly represented adult-stage cells.

We further validated our cluster annotation by evaluating the enrichment of cluster-specific differentially expressed genes (DEGs) with cell type-specific DEGs identified in the latest single-cell study of the developing human brain^20^ (**Figure 1H**; **Table S2**). We observed a significant overlap between fetal excitatory neurons (C12, C29) and previously identified early fetal excitatory neurons (EN-fetal-early) (C12, odds ratio [OR] = 7.81, false discovery rate [FDR] = 9.98 x 10^-21^; C29, OR = 6.35, FDR = 3.08 x 10^-20^), as well as late fetal excitatory neurons (EN-fetal-late) (C12, OR = 8.15, FDR = 1.51 x 10^-29^; C29, OR = 7.55, FDR = 9.57 x 10^-36^). We found a significant overlap between fetal inhibitory neurons (C20, C23) and previously identified fetal inhibitory neurons (IN-fetal) (C20, OR = 8.38, FDR = 1.25 x 10^-9^; C23, OR = 6.70, FDR = 8.52 x 10^-6^). Radial glia, neuroblasts, and non-neuronal cell types also exhibited significant enrichment with the cell-type-specific DEGs, further confirming our classification results. Regarding C38, which predominantly consists of oligodendrocytes present during the neonatal stage, significant overlap was observed with both oligodendrocytes (OR = 6.25, FDR = 1.25 x 10^-2^) and OPC (OR = 28.8, FDR = 3.15 x 10^-33^), suggesting an ongoing differentiation process towards mature oligodendrocytes within this cluster. Overall, these results validate the annotation and underscore the robustness and fidelity of our atlas in capturing the dynamic landscape of human brain development.

### Cellular landscape of neurodevelopmental disorder genes in early neuronal lineages

Over the past decade, large-scale genomic studies have identified risk genes associated with neurological disorders and implicated a substantial locus heterogeneity in the underlying etiology. Unraveling the intricate temporal patterns of risk genes is crucial for deciphering the underlying pathological mechanisms and identifying the most relevant cell types and developmental stages implicated in disease pathogenesis. To investigate the cellular and temporal specificity, we compared the risk genes associated with these disorders with genes enriched in specific cell subtypes (**Figure 2A**; **Table S3A**). We have prioritized 14 sets of genes previously reported to be associated with neurological disorders in large-scale genomic studies and examined their expression profiles in our single-cell atlas. Consequently, distinct patterns of gene enrichment were detected in a cell type-specific manner. Risk genes for neurodevelopmental disorders, including autism, developmental delay, and epilepsy, are predominantly expressed in neuronal cell types. Autism and developmental delay risk genes were considerably enriched in both excitatory and inhibitory neuronal cell types, which is consistent with previous findings^21^ (**Table S3B**). Genes associated with epilepsy demonstrate specific enrichment in excitatory neurons located in cortical layers 4-6 (C36) and SST-expressing inhibitory neurons (C6).

**Figure 2.**
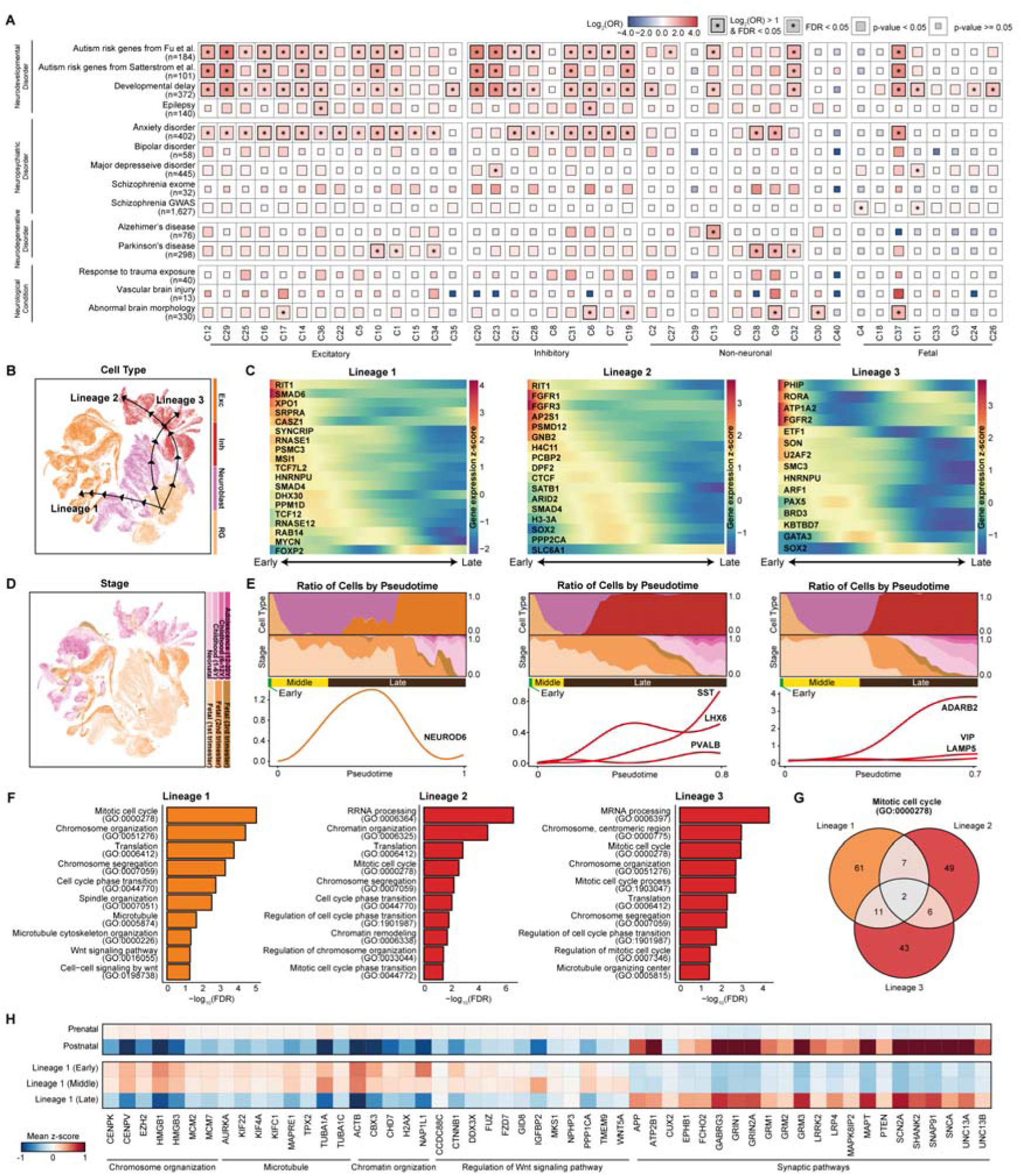
Cellular trajectories of neurodevelopmental disorder risk genes in neuronal lineages. **A.** Gene set enrichment test for neurological disorder genes. A one-sided Fisher’s exact test was used to compute statistics with multiple comparisons by FDR. **B.** UMAP visualizations of estimated developmental lineages in neuronal cell types. **C.** Expression profiles of neurodevelopmental disorder risk genes across pseudotime for each lineage. **D.** UMAP visualizations of developmental stages in neuronal cell types. **E.** Distribution of cells by major cell type and developmental stage across pseudotime and temporal patterns of late neuronal IPC marker (NEUROD6), MGE-derived inhibitory neuron markers (SST, LHX6, and PVALB), and CGE-derived inhibitory neuron markers (ADARB2, VIP, and LAMP5). Cells in the first one-third portion of cells with pseudotime close to 0 are labeled as "Early", the subsequent portion as "Middle", and the final one-third of cells with the latest pseudotime are designated as "Late". **F.** Functional annotations for lineages with significantly enriched biological processes with multiple comparisons by FDR. **G.** Venn diagram illustrating the number of overlapped genes involved in the mitotic cell cycle pathway across lineages. **H.** Heatmap depicting z-score normalized expression of genes involved in lineage timepoint-specific biological pathways. Synaptic pathways include genes involved in chemical synaptic transmission (GO:0007268) and synaptic membrane (GO:0097060).

We further investigated the cellular lineages underlying neuronal clusters and the lineage-specific patterns of neurodevelopmental disorder risk genes. The pseudo-time trajectory built on cellular identity reflected the clustering patterns of cells differentiating from radial glia to neuroblasts, eventually committing to mature neurons under the age of 20 years (**Figure S2**). Three distinct lineages diverged from radial glia and differentiated into excitatory and inhibitory neurons (**Figure 2B**), each enriched for specific risk genes (**Figure 2C**) and encompassing cells across various developmental stages (**Figure 2D**). These lineages were characterized as those destined for excitatory neurons (lineage 1), MGE-derived inhibitory neurons (lineage 2), and CGE-derived inhibitory neurons (lineage 3) (**Figure 2E**).

Lineage 1 represents a differentiation trajectory from radial glia to excitatory neurons, marked by the transient expression of NEUROD6 as neuroblasts transition into excitatory neurons. Among the autism risk genes, those involved in transcriptional regulation, such as RNASE1 and TCF7L2, are predominantly expressed in the early stages of this lineage. Most autism and developmental delay risk genes peaked during the initial phase of the lineage. In contrast, RAB14 and MYCN, other risk genes for developmental delay, exhibited peaks in later stages, coinciding with neuroblast differentiation into excitatory neurons during the fetal second trimester. FOXP2, a risk gene for both autism and developmental delay, showed a burst expression at the end of the lineage, particularly in the postnatal stage. The Wnt signaling pathway (GO:0016055) mediated by TCF7L2 was significantly enriched in this lineage (FDR = 4.79 x 10^-2^) (**Figure 2F; Table S4**), suggesting its involvement in the pathophysiology of autism and developmental delay. Lineage 1 is also characterized by enrichment in cell cycle and translation processes, implying heightened cellular activity and proliferation, which are essential for neurogenesis and neuronal differentiation. Though the enrichment of mitotic cell cycles (GO:0000278) was recurrent in other lineages, the low number of overlapped genes confirms the lineage-specific characteristics distinct from other lineages (**Figure 2G**).

To further elucidate the distinctions among the pseudo-time stages within this differentiation lineage, we compared cells at earlier, middle, and later pseudo-time points along with their enriched biological pathways (**Figure 2H**). Cells transitioning from early to middle pseudo-time stages predominantly exhibited enrichment in pathways related to cellular infrastructure, such as chromosome organization (Early, FDR = 1.74 x 10^-5^; Middle, FDR = 2.26 x 10^-8^) and microtubule-based process (Early, FDR = 7.17 x 10^-3^; Middle, FDR = 3.61 x 10^-2^). Genes involved in these pathways, including centromere protein (CENPK and CENPV), high mobility group box (HMGB1 and HMGB3), and tubulin alpha (TUBA1A and TUBA1C), are highly expressed in early- and middle-stage cells than in later stages. Nonetheless, cells in the middle pseudo-time stage were also specifically enriched in the regulation of the Wnt signaling pathway (FDR = 4.15 x 10^-2^), mediated by up-regulation of genes such as FZD7, GID8, IGFBP2, TMEM9, and WNT5A. Cells in the later pseudo-time stage exhibited specific enrichment in synaptic pathways, characterized by the upregulation of several neuronal markers including CUX2, glutamate metabotropic receptor genes (GRM1, GRM2, and GRM3), and several autism risk genes including PTEN, SCN2A, and SHANK2. These results indicate that essential biological processes are dynamically regulated throughout neurodevelopment, highlighting the temporal specificity of the differentiation.

Lineages 2 and 3 represented the differentiation trajectory from radial glia to inhibitory neurons, diverging into MGE-derived (C20) and CGE-derived (C23) neurons (**Figure S2**). This divergence is distinguished by the increased expression of the MGE-branch markers SST, LHX6, and PVALB in lineage 2, and the increased expression of CGE-branch markers ADARB2, VIP, and LAMP5 in lineage 3. In lineage 2, regarding autism risk genes involved in neuronal communication, AP2S1 exhibited initial expression, whereas SLC6A1 was highly expressed in a later lineage. Risk genes for developmental delay, including CTCF, SATB1, ARID2, SMAD4, H3-3A, SOX2, and PPP2CA, are coherently expressed during the transition phase from neuroblasts to fetal inhibitory neurons around the fetal second trimester, suggesting a heightened risk of disorder at this stage of neural development. In lineage 3, risk genes associated with developmental delay, including PHIP, RORA, ATP1A2, FGFR2, ETF1, SON, U2AF2, SMC3, and HNRNPU were coherently peaked during the initial phase of the lineage. PAX5, which represents an autism risk gene involved in gene expression regulation, peaks during the middle phase of this lineage, within neuroblast cells of the fetal first trimester. Moreover, other risk genes for developmental delay, such as BRD3, KBTBD7, and GATA3, also showed peaked during this stage, whereas SOX2 displayed high expression in a later lineage. Coherent to lineage 1, lineages 2 and 3 exhibited significant enrichment in the pathways associated with chromosomal organization and mitotic cell cycle processes. Nonetheless, few genes overlapped between the common pathways, implying that lineage-specific genes distinctly constitute the core pathways in early neuronal development. These results suggest that the distinct association of risk genes with neuronal lineage aligns with functional variations across neuronal maturation.

### Exploring the neurological disorder gene expressions in glial cell types

Glial cells play a pivotal role in maintaining nervous system homeostasis, providing support and protection to neurons, and participating in signal transmission. The dysfunction of glial cell types has been reported in several neurodegenerative disorders, such as Alzheimer’s disease and Parkinson’s disease. Nonetheless, the detailed trajectories of glial cell differentiation and their implications in neurological disorders remain unknown. As described previously, we mapped the risk genes for neurological disorders in a developmental trajectory of glial cell types (**Figure 2A**). Astrocytes (C2, C27) exhibited significant enrichment of risk genes for developmental delay (C2, OR = 2.22, FDR = 1.69 x 10-4) and autism (C27, OR = 1.96, FDR = 3.67 x 10^-3^). Risk genes of Alzheimer’s disease were predominantly expressed in microglia (C13, OR = 3.83, FDR = 3.96 x 10^-3^), emphasizing its potential role in the pathology or progression of Alzheimer’s disease. The risk genes for Parkinson’s disease demonstrated were specific enrichment in fetal oligodendrocytes (C38) (OR = 3.22, FDR = 2.53 x 10^-4^) and OPCs (C9, OR = 2.86, FDR = 5.65 x 10^-6^; C32, OR = 1.89, FDR = 2.92 x 10^-3^).

To further investigate the expression dynamics of these risk genes, we identified individual cellular trajectories for each cell type and the expression profiles of the risk genes across them. Oligodendrocytes appear to have a single trajectory, which starts from OPCs (C9 and C32) and differentiates into fetal oligodendrocytes (C38) and mature oligodendrocytes (C0) (**Figure 3A**). This trajectory was characterized by mixed developmental stages across the lineage pseudo-time, suggesting the presence of both progenitor and mature cells throughout the postnatal period (**Figure 3B, 3C, S3**). Risk genes specifically expressed in the oligodendrocyte lineage exhibited distinct peak expression patterns (**Figure 3D**). In the earlier lineage, risk genes associated with abnormal brain morphology (AGMO) and anxiety disorders (KAT2B) showed high expressions. Earlier genes were involved in neuronal projections and intracellular cytoskeletal activity (**Figure 3E**). This included a microtubule-based process (GO:000701, FDR = 1.96 x 10^-2^) mediated by KAT2B, a risk gene for abnormal brain morphology. SOX10, a risk gene associated with developmental delay, showed the highest expression at the stage when OPCs differentiated into oligodendrocytes. Conversely, PHLDB1 spikes as oligodendrocytes mature, implying temporal variation in the expression of developmental delay risk genes during oligodendrocyte development. Two risk genes of Parkinson’s showed distinct patterns, with PLPP4 expressed in initial OPC cells and DNAH17 expressed in the late oligodendrocytes. Other genes expressed in the latter part of the lineage include anxiety disorder risk genes (AATK and TBC1D2), abnormal brain morphology genes (TSPAN15, GLTP, and SLCOB1), and an autism risk gene (ATG13). Cells at a later stage exhibited enrichment in pathways related to cell growth and morphogenesis (**Figure 3E**), suggesting a maturation process rather than rapid differentiation and transition of cell state.

**Figure 3.**
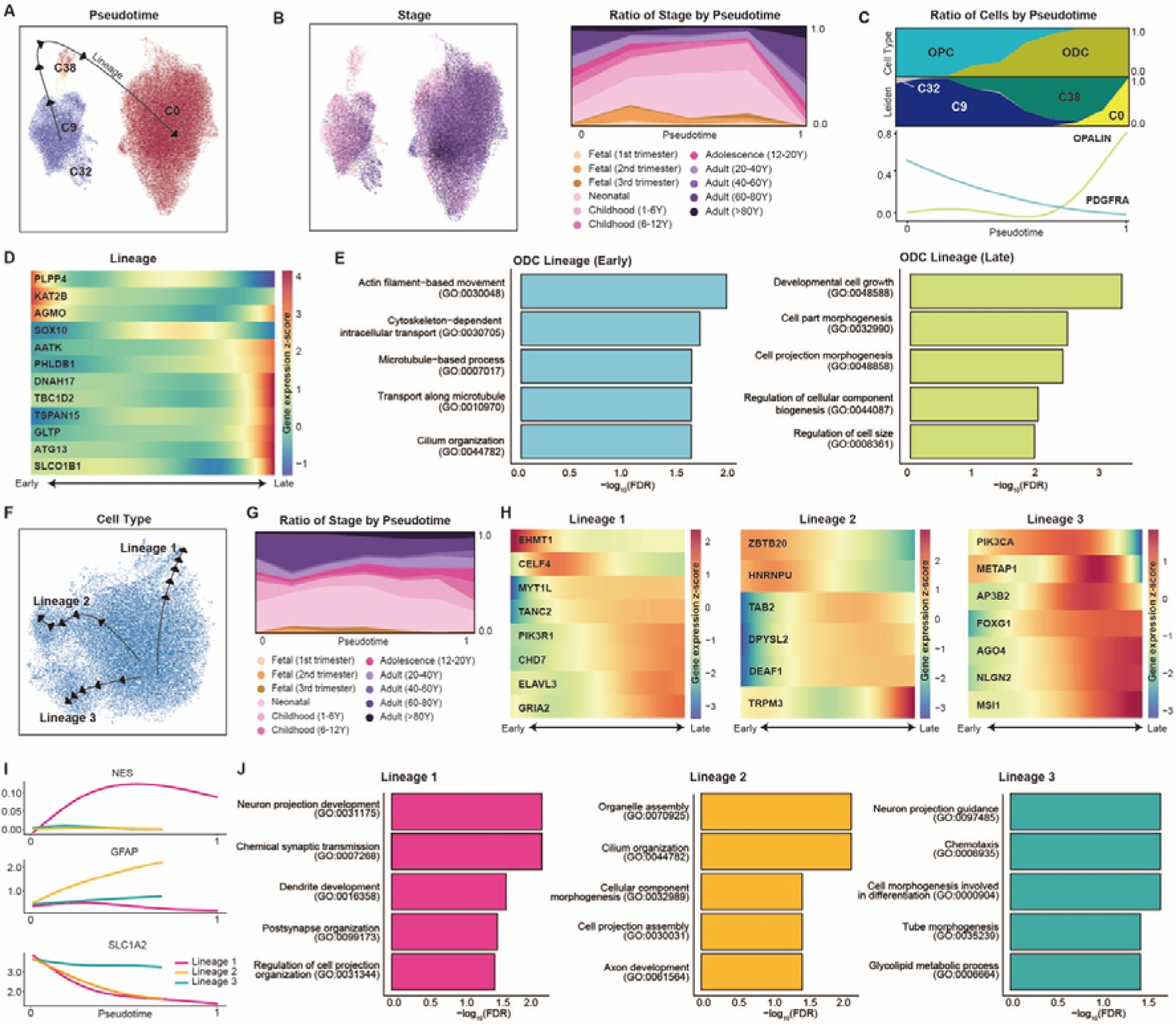
Cellular trajectories of neurological disorder risk genes in non-neuronal lineages. **A-B.** UMAP visualization of estimated developmental lineage in oligodendrocyte group (C0, C8, C9, and C38), colored by pseudotime (A) and developmental stage (B). **C.** Distribution of cells by major cell type and cluster across pseudotime and temporal patterns of PDGFRA and OPALIN. **D.** Expression profiles of neurological disorder risk genes across pseudotime for lineage. **E.** Functional annotations for early and late lineage cells with significantly enriched biological processes with multiple comparisons by FDR. **F.** UMAP visualization of the estimated developmental lineage in the astrocyte group (C2 and C29). **G.** Distribution of cells colored by developmental stage across pseudotime. **H.** Expression profiles of neurological disorder risk genes across pseudotime for each lineage. **I.** Distinct expression pattern of known reactive astrocyte marker genes in each lineage. **J.** Functional annotations for each lineage with significantly enriched biological processes with multiple comparisons by FDR.

Astrocytes possess three distinct lineages, regardless of the developmental stage or clusters (**Figure 3F–J, S3E–I**). These lineages were further characterized by the distinct expression of risk genes and known markers involved in diverse functions of reactive phenotypes^33^. In lineage 1, the autism risk gene EHMT1 was highly expressed at the earlier point, followed by the sequential peaking of CELF4. Risk genes of developmental disorders (ELAVL3, GRIA2, and CHD7) exhibited a high expression at the later stages of the lineage. This lineage is enriched in neuron projections and synaptic transmission pathways. Lineage 1 was exclusively enriched in a type of intermediate filament (IF) gene nestin (NES), which is also a marker of neural stem and progenitor cells. Pathways enriched in this lineage are mainly associated with neuronal development, implying its neurodevelopmental role. In lineage 2, risk genes of developmental delay, such as ZBTB20 and HNRNPU, were highly expressed at the initial stage. In contrast, TAB2 expression peaked during the middle stage, whereas TRPM3 exhibited strong expression at the latter stage, suggesting distinct roles for each gene during cell maturation within the lineage. Autism risk genes involved in transcriptional regulations (DEAF1) and cytoskeleton (DPYSL2) were strongly expressed at the end of the lineage. This lineage is characterized by distinct expression of GFAP along the progress of maturation, which is another well-known gene of IF as well as a hallmark for reactive astrocyte phenotypes^34^. Cilium organization pathway (GO:0044782, FDR = 7.68 x 10^-2^) is also enriched in this lineage, where the cilium serves as a microtubule-based signaling device for various physiological functions of astrocytes. In lineage 3, PIK3CA, a risk gene for both autism and developmental delay, peaked at the middle part of the lineage. This is followed by the sequential expression of METAP1, a risk gene for autism and schizophrenia, along with AP3B2, a risk gene for developmental delay. The expression levels of other genes associated with developmental delay (FOXG2, NLGN2, and MSI1) peaked later in the lineage. Lineage 3 is also characterized by constant expression of genes related to glutamate transporters. Overall, these results suggest an association of risk genes with cellular maturation lineages in non-neuronal cells, indicating a temporal specificity of disorder risk.

### Multifaceted regulatory mechanisms governing early brain development

To deepen our understanding of the complex orchestration of brain development, we investigated the regulatory landscape of transcription factors, immune signaling, hormonal regulation, and kinase-mediated pathways. First, we predicted a cluster-specific gene regulatory network linking transcription factors to their putative target genes by integrating motif enrichment and gene co-expression analyses (**Table S5A**). In total, 633 transcription factors were found to activate in at least one cell type (**Figure 4A**). Based on the predicted regulon activity of the transcription factors, the clusters were categorized into six different groups. From this, we identified canonical transcription factors for neurodevelopment, such as NFIA (astrocytes)^35,36^ and TCF12 (oligodendrocytes)^37^, confirming the validity of our findings.

**Figure 4.**
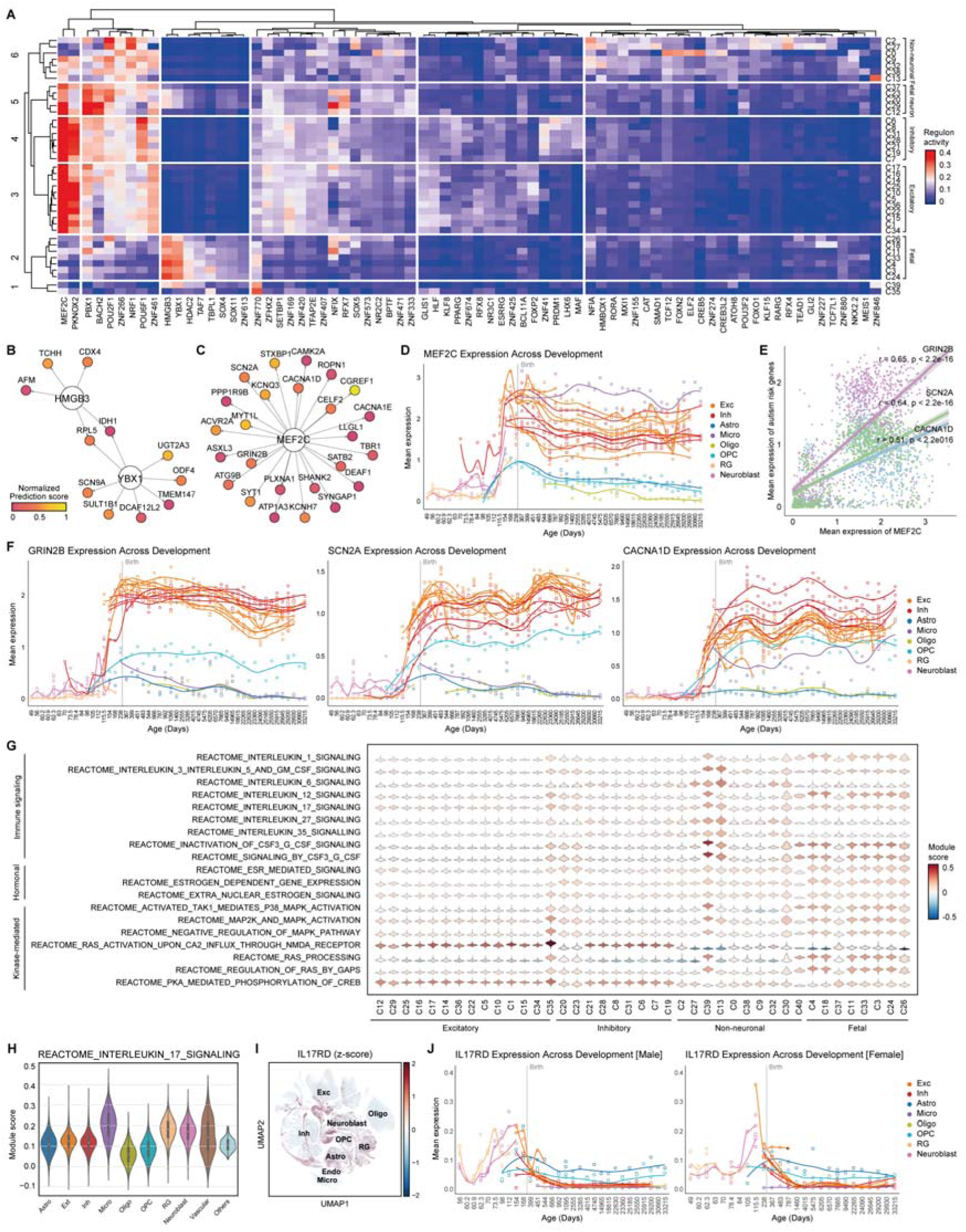
Regulatory landscape and pathway enrichment in early brain development. **A.** Heatmap illustrating regulon activities of transcription factors across clusters. **B–C.** Transcription factor-target networks depicting regulation of glioblastoma risk genes by HMGB3 and YBX1 (B), and regulation of autism risk genes by MEF2C (C). Prediction confidence was normalized from 0 to 1. The top 25 targets high-confidence targets for MEF2C are shown. **D.** Expression of MEF2C over gestational days. The sample-wise mean of log-normalized MEF2C expression was computed using a pseudo-bulk method. Clusters with at least 4,600 cells (C0-C22) were used. **E.** Correlation between the sample-wise mean of log-normalized MEF2C expression and expression of GRIN2B, SCN2A, and CACNA1D. **F.** Expression of GRIN2B, SCN2A, and CACNA1D across developmental ages. **G.** Violin plot displaying pathway module score as the average expression level of pathway genes adjusted for control features. **H.** Violin plot for pathway module score across major cell types. **I.** UMAP visualization of z-score normalized IL17RD expression. **J.** Expression of IL17RD over gestational days, divided by gender.

Based on these results, we further elucidated the transcription factors implicated in neurological disorders. Fetal radial glia and neuroblast cells were enriched for HMGB3, and YBX1 (Group 2). These genes are involved in glioblastoma tumorigenesis and tumor growth^38, 39^. We found that HMGB3 and YBX1 are predicted to target 11 risk genes for glioblastoma, including RPL5 and IDH1, as shared target genes (**Figure 4B**). Fetal neurons appear to have different transcription factors for later differentiation from radial glia and neuroblasts. Fetal neurons are regulated by MEF2C, PBX1, BACH2, POU2F1, NFIX, and RFX7 (Group 3). NFIX, known for its involvement in the differentiation of radial glia into neuronal IPC^40^, and RFX7, known for its role in neural tube information^41^, exhibited distinct activities within this group. NFIX has been predicted to target 28 autism risk genes, including DIP2C and CSNK1E. MEF2C, which further displays potent activity in adult neurons, is a transcription factor that regulates multiple genes associated with autism^42^. We identified 47 autism risk genes that were predicted to be targeted by MEF2C (**Figure 4C**). The expression level of MEF2C increased in neurons across developmental stages, particularly in the fetal third trimester (**Figure 4D**). This pattern was consistent with that of other autism risk genes, such as GRIN2B (correlation coefficient (r) = 0.65, p < 2.2e-16), SCN2A (r = 0.64, p < 2.2e-16), and CACNA1D (r = 0.51, p < 2.2e-16), suggesting a coordinated increase during the fetal third trimester (**Figures 4E, 4F**). These results suggest that the activation of MEF2C plays a leading role in controlling the expression of autism risk genes.

We further compared the signature genes of hormonal regulation, kinase-mediated, and immune signaling pathways across cell types to elucidate their contributions to neurodevelopment (**Figure 4G, S4**; **Table S5B**). Pathways associated with the sex hormone, estrogen, showed greater enrichment in fetal neurons compared to adult neurons, implying differential responsiveness to hormonal cues during neural maturation. Conversely, kinase activity, especially within the RAS pathways, was predominantly enriched in adult neurons compared to fetal neurons, implying its role in neuronal survival and regeneration rather than in neurodevelopment.

Immune signaling pathways were highly enriched in fetal and adult microglia compared with other cell types (**Table S5C**), indicating their pivotal role in immune activities within the brain. Along with microglia, radial glia and neuroblasts showed overexpression of immune pathways, particularly interleukin (IL) signaling pathways (**Figure 4H**). Previous studies have reported a putative role for IL-6 and IL-17A in the development of autism-like behavioral phenotype in mice models of maternal immune activation (MIA)^43,44^. Autism-like behavior in the MIA mouse model was mediated by IL-17 receptor expression in the offspring brain^45^. While some of the IL-17 receptor genes (IL17RA, IL17RB, IL17RC, and IL17RE) were not distinctly expressed in the human fetal brain (**Figure S5**), IL17RD was elevated in fetal radial glia and neuroblasts (**Figure 4I**), implicating its role in a prenatal risk to MIA. We further examined the onset of IL17RD expression and found sex-dependent differences (**Figure 4J**). In males, IL17RD expression in radial glia and neuroblasts begins around gestational day 60, peaks between days 112 and 154, and then decreases postnatally. In females, IL17RD expression begins later, rising rapidly between gestational days 84 and 105, peaking around day 115.5, and decreasing after birth. As MIA offspring were known to exhibit male-biased behavioral abnormalities^46^, these findings may implicate that the differential regulation of IL17RD expression between males and females may contribute to the varying susceptibility to MIA.

### Identification of cellular characteristics underlying glioblastoma

Glioblastoma is a lethal primary brain tumor characterized by intra-tumoral heterogeneity. A single-cell atlas may help in understanding the cellular heterogeneity underlying the clinical and molecular complexity of this disorder. We first examined whether the glioblastoma driver genes^26^ overlapped with cell type-specific genes. None of the cell types showed significant enrichment. This suggests that genomic associations might not fully characterize the molecular underpinnings of glioblastoma.

Thus, we examined the cellular specificity of the signature gene sets, previously defined to represent diverse cellular states in glioblastoma^27^ (**Figure 5**). Overall, the signature gene sets were aligned with the corresponding original cell types. Neural progenitor cell (NPC)-like tumors were significantly enriched in fetal neurons and fetal cell types. NPC-like subprogram 1 cell genes showed enrichment for OPC and oligodendrocyte cells. Among the signature genes, BCAN, DLL3, OLIG1, LRRN1, TCF12, and TNR were expressed specifically in these cell types. In contrast, NPC-like subprogram 2 cell genes demonstrated significant overlap for layers 4-6 excitatory neuron (C17) and layer-unspecific excitatory neuron (C35). TCF4 was included in the NPC-like subprogram 2 cell genes, emphasizing its regulatory function in tumor progression.

**Figure 5.**
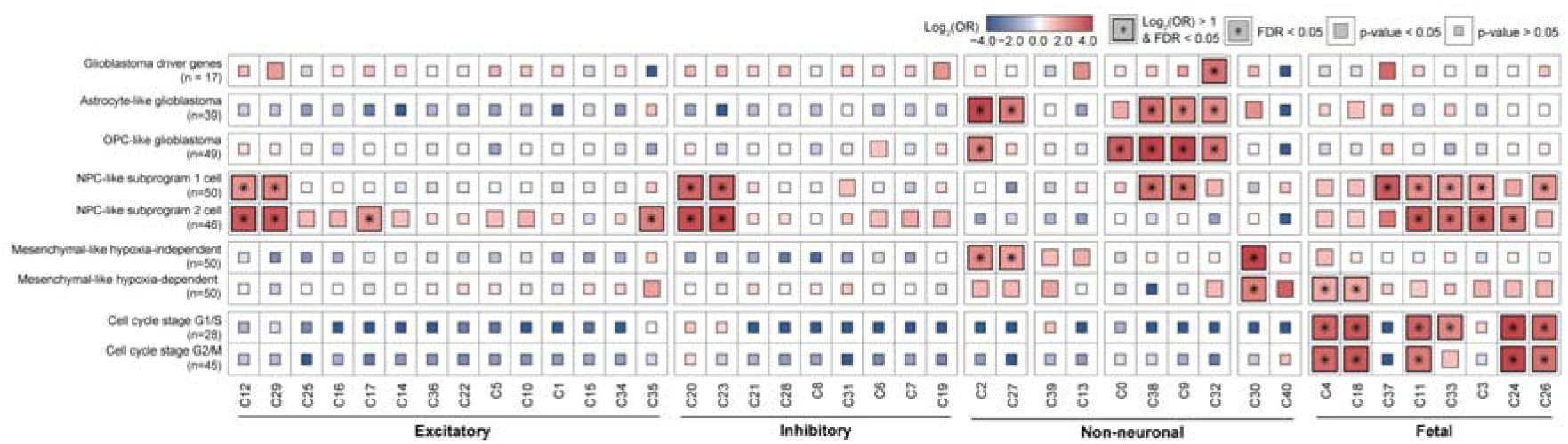
Cell type and temporal specificity in glioblastoma subtypes. Gene set enrichment test with driver genes and transcriptional signatures of glioblastoma. A one-sided Fisher’s exact test was used to compute statistics with multiple comparisons by FDR.

Mesenchymal-like tumor-related genes exhibited different enrichment patterns between hypoxia-independent and hypoxia-dependent subgroups. While both were enriched in vascular cells (C30), hypoxia-independent tumor-associated genes were enriched in astrocytes (C2, C27), and hypoxia-dependent tumor-related genes showed enrichment in radial glia subtypes expressing HES1 (C4, C18), implying a possible resemblance of each astrocyte and radial glia subtype to mesenchymal-like malignant cells. Furthermore, genes containing information on the cell cycle phases G1/S and G2/M were significantly enriched in fetal cells, implying the importance of studying fetal cells even in non-fetal or neonatal cases. These results highlight the significance of investigating cell type-specific perspectives and studying brain cell types across primary developmental stages to gain better clinical insights into disease initiation and potential treatment options.

## Discussion

In this study, we constructed a comprehensive single-cell atlas of the developing human brain to investigate the cellular and temporal specificity of genes implicated in neurological disorders. An atlas-level integration enables detailed functional and molecular profiling of various cell types across multiple human samples. By analyzing 393,060 single brain cells, our research revealed the complex cellular composition and dynamic changes during early brain development.

Neurological disorders are characterized by a high level of genetic heterogeneity, wherein multiple genes may be associated with a single disorder. Despite the huge success of large-scale genomic studies, such as GWAS, these associations require appropriate interpretation in a biological and genomic context. Consequently, deciphering the functional convergence of risk genes is essential for comprehending disease pathophysiology and identifying potential therapeutic targets. Our single-cell atlas can be useful for thorough analysis of risk genes associated with neurological disorders that may occur during brain development. We found the distinct expression pattern of the autism risk gene FOXP2, which highlights its temporal role in excitatory and inhibitory neuronal lineages and offers insights into its developmental implications. PD risk genes, PLPP4 and DNAH17, were shown to have temporal regulation at distinct time points during oligodendrocyte differentiation, implying temporal specificity appears not only in neuronal cells but also in glial cells. Moreover, the atlas enabled the exploration of novel disease mechanisms. For example, it elucidates the role of MEF2C, a transcription factor activated during the transition from prenatal to postnatal neurons, in alignment with the expression patterns of autism risk genes. In addition, the distinct sex difference in the onset of IL17RD expression may indicate it as a putative target contributing to varying susceptibility to MIA. Furthermore, we revealed that certain cellular states in glioblastoma closely resemble fetal-stage cell types despite their origin in adults, implying the necessity to investigate the characteristics leading to this similarity.

This discovery comprehensively delineates the regulatory mechanisms in early brain development and facilitates advancements in research and potential therapeutic interventions. There has been extensive research focused on studying the cell-type-specific nature of neuronal diseases, primarily focusing on stage-specific neuronal cells. However, to enhance clinical investigations and therapeutics further, it is crucial to consider the temporal and cellular specificity of neuronal disorders, as well as for the non-neuronal cell types, which also exhibited temporal specificity across cellular maturation. Moreover, our study of neuronal disorders indicates a significant enrichment in fetal cell types. This suggests that the determination of disorders may be strongly influenced during the prenatal stage. Therefore, we propose that studying temporal specificity across brain development warrants a more robust investigation.

Although our single-cell atlas leveraged a large number of postmortem samples to explore the developing human brains, our study had several limitations. Despite using data from various donors, our atlas may not encompass the full spectrum of individual variability in developmental trajectories. Considering the pivotal role assumed by the genetic constitution in early brain development and its impact on gene expression^5^, a more extensive and diverse collection of samples is required to enhance our understanding. Furthermore, integrating single-cell ATAC sequencing data could provide valuable insights into regulatory enhancers and mechanisms of brain development. Another limitation is posed by the scarcity of samples from the neonatal and early childhood periods. These stages are crucial for assessing synaptic pruning and the maturation of neural circuits; nevertheless, the availability of human postmortem brain samples from these stages is constrained. Addressing this gap warrants concerted efforts to collect and analyze samples from these critical developmental stages to enrich comprehension regarding brain development and its implications for neurological disorders.

## Data availability

The weights from the scVI model for reference mapping with scArches, integrated embedding, and metadata are publicly available on Zenodo. Plots illustrating the expression profile for 3,380 neurological disorder risk genes across the atlas are also provided. Further availability for the data can be inquired to the corresponding author.

## Acknowledgment

This study was supported by the National Research Foundation of Korea (NRF) grant funded by the Korea government (NRF-2020R1C1C1003426 and NRF-2022M3E5E8018049 to J.Y.A.) the Korea Health Technology R&D Project through the Korea Health Industry Development Institute and Korea Dementia Research Center, funded by the Ministry of Health & Welfare and Ministry of Science and ICT, Republic of Korea (grant number: HU22C0042), and Korea University to J.Y.A. S.Y.K. received from the Brain Korea (BK21) FOUR education program.

## Author contributions

Study design: S.K. and J.Y.A. Data Collection: S.K., I.G.K, J.L., and J.Y.A. Data analysis: S.K., J.L., I.G.K, J.J and J.Y.A. Manuscript Writing: S.K., J.L., and J.Y.A. Review and Feedback: E.H.K., H.J.K., J.P., J.E.P., and J.Y.A. All authors have read and approved the final version of the manuscript for publication.

---

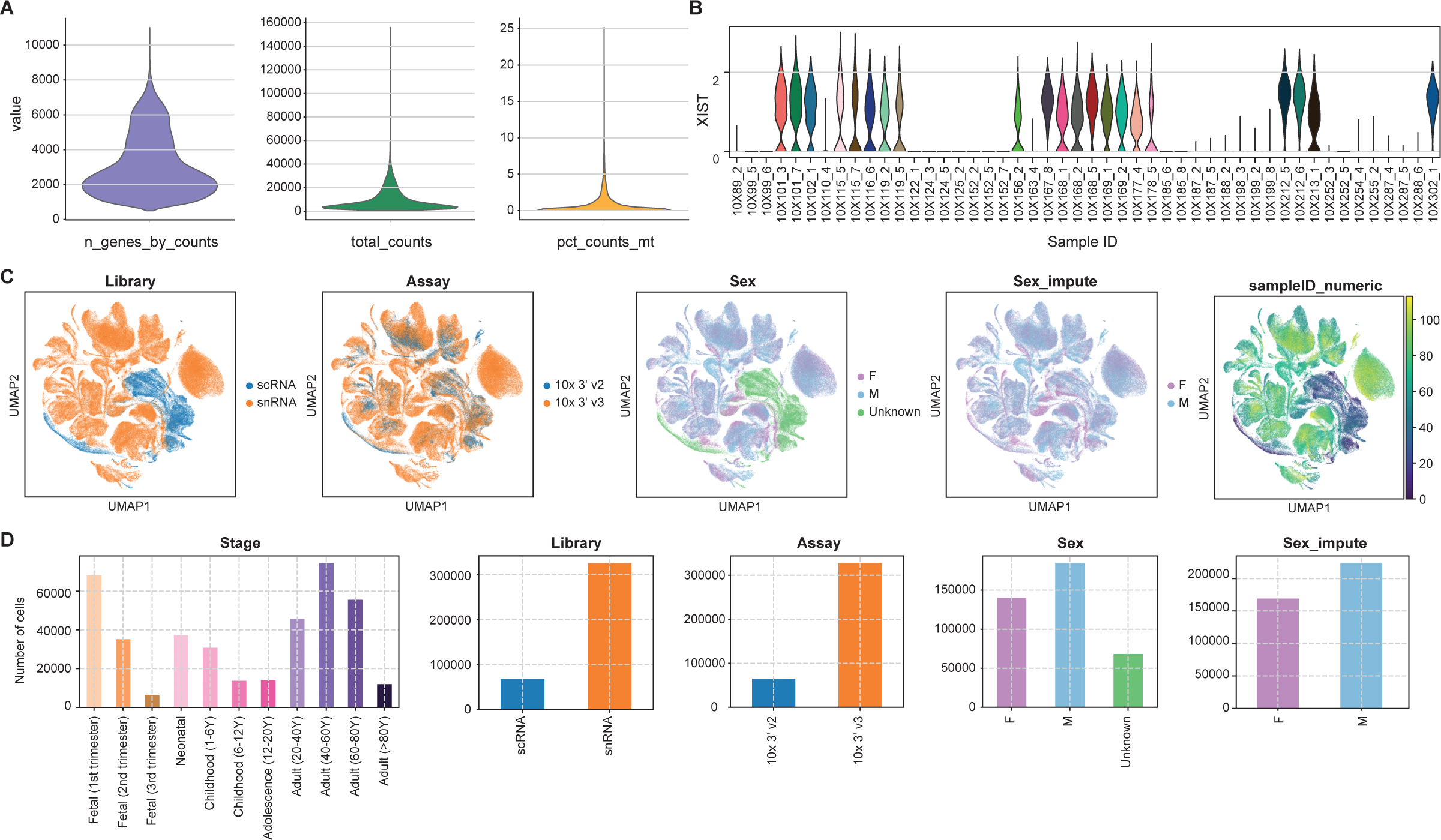

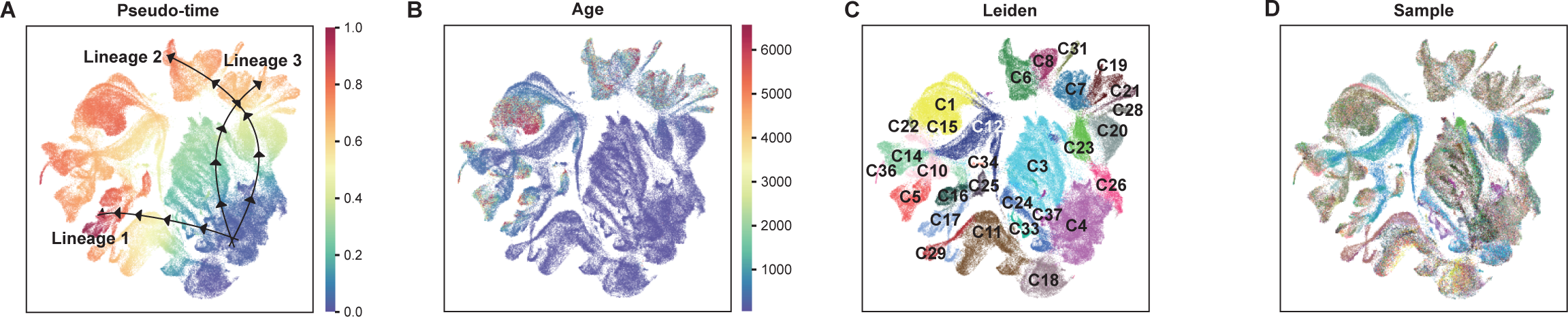

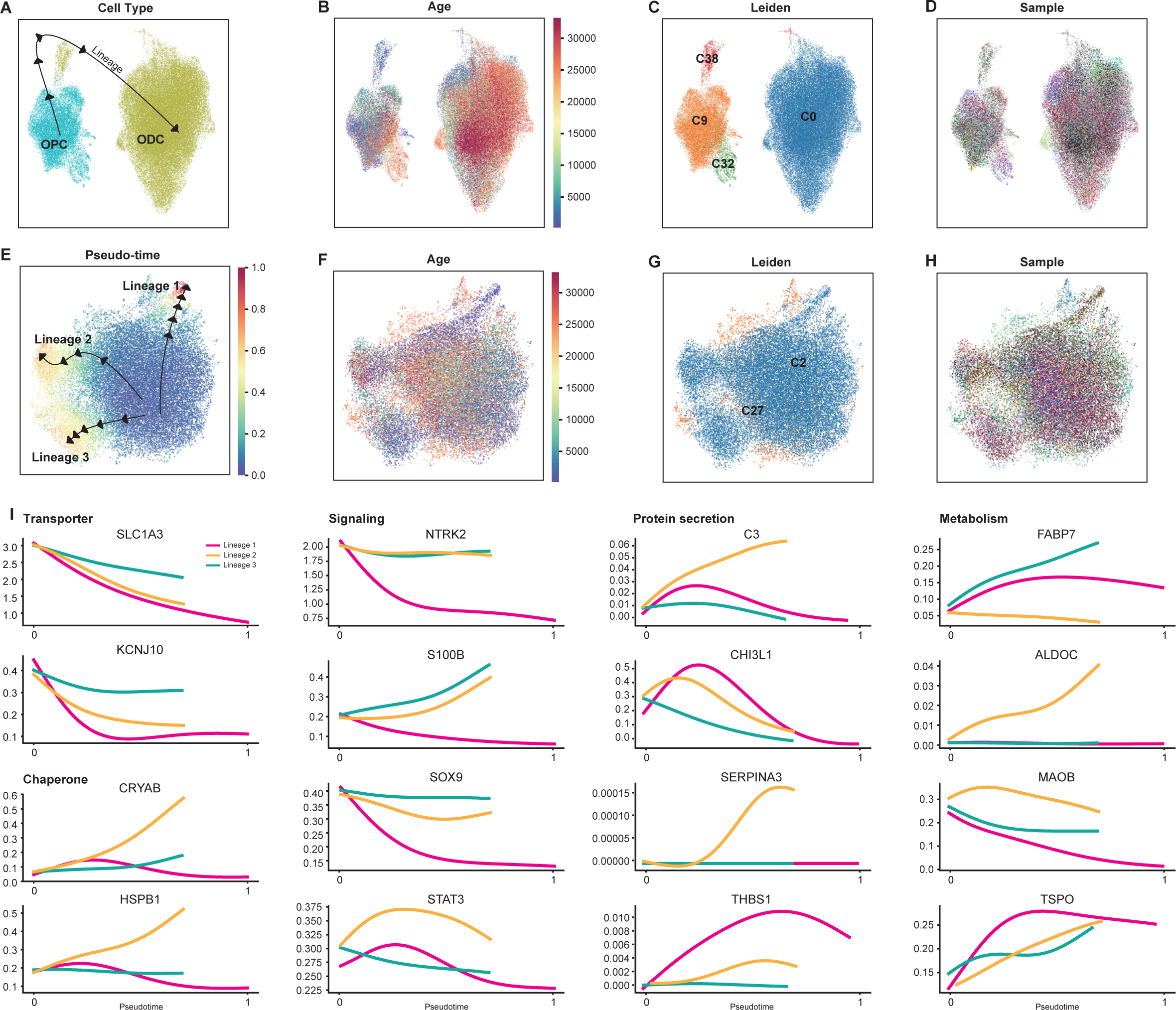

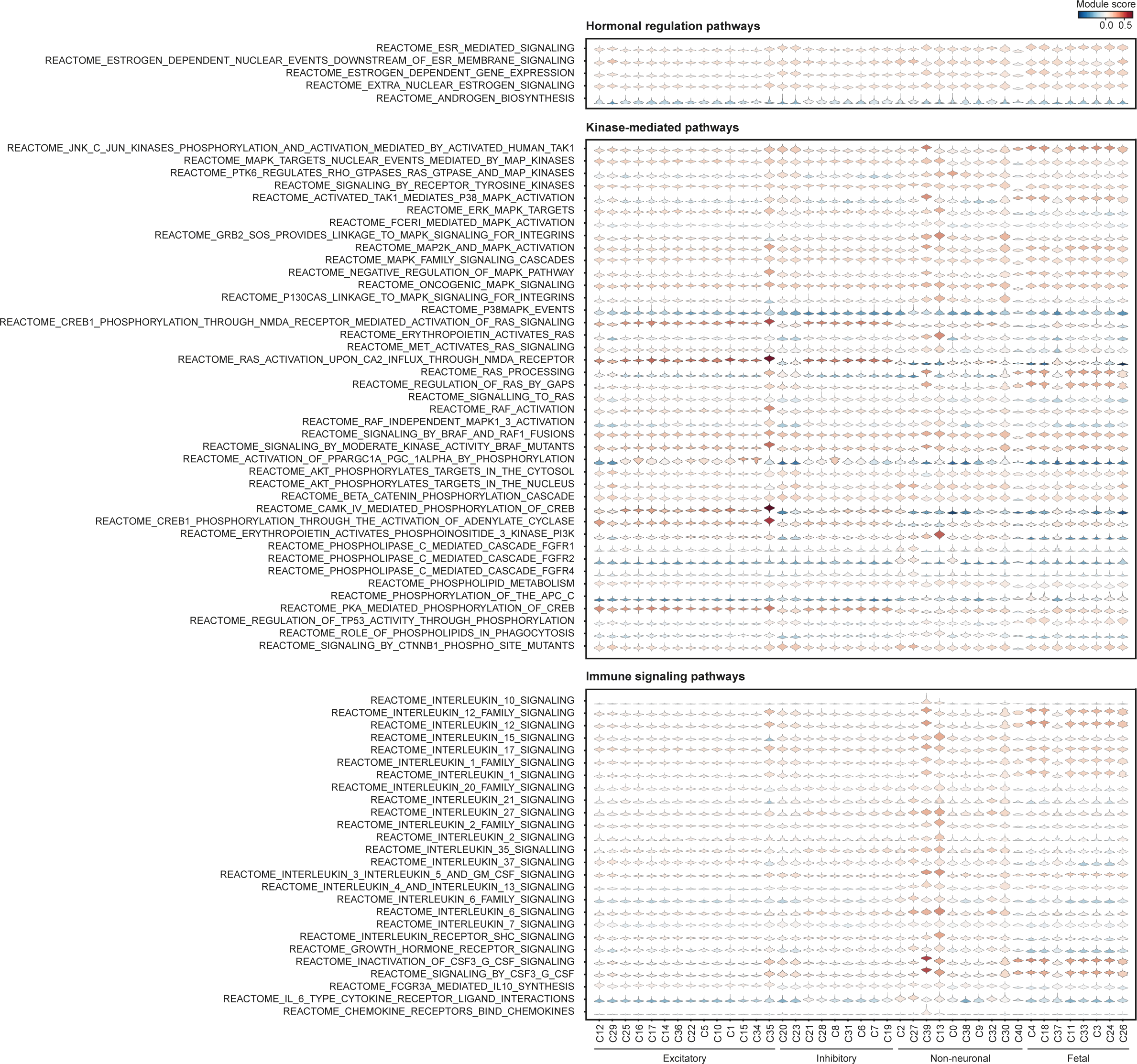

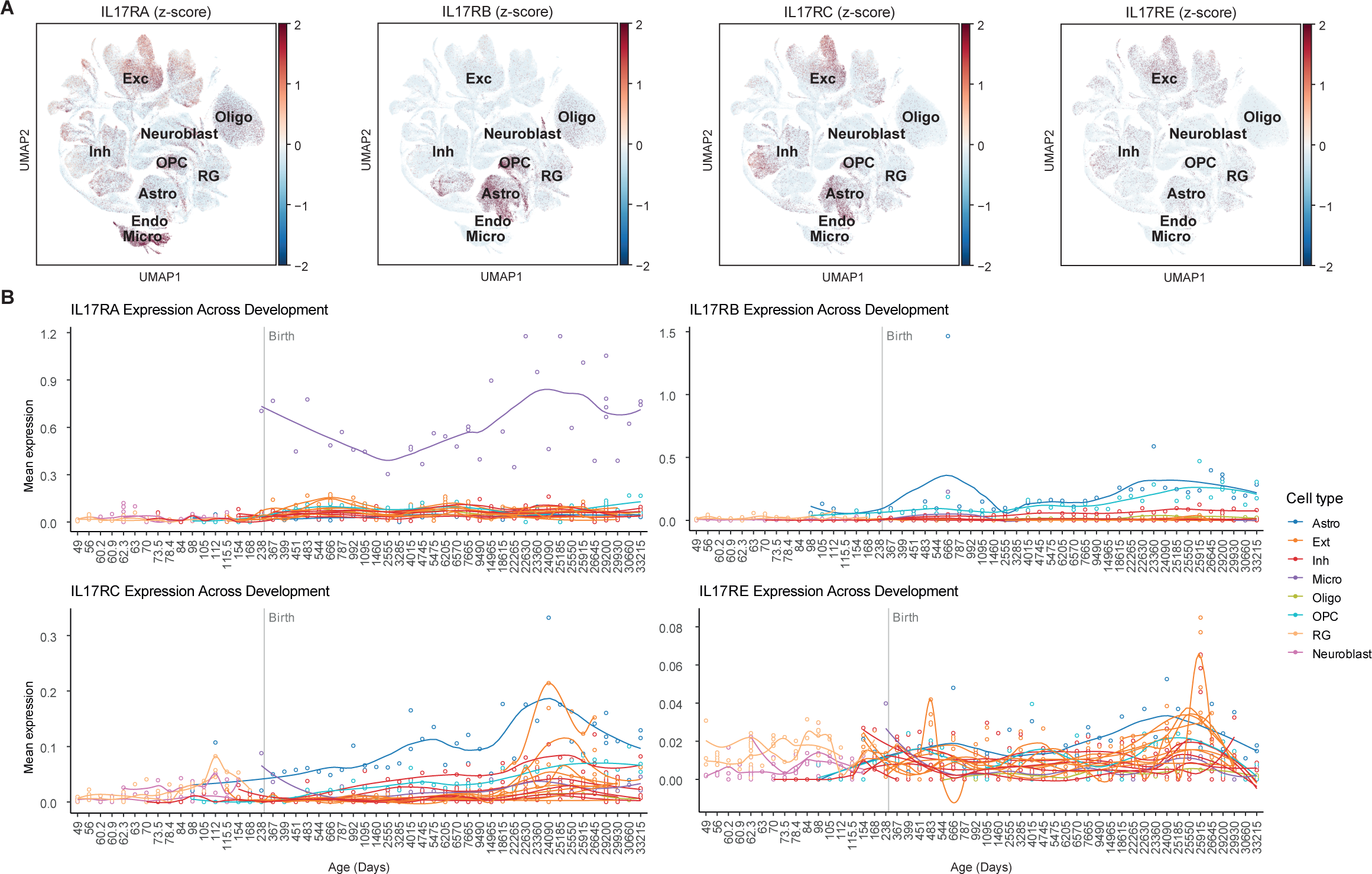

## References

1 Bakken, T. E. et al. Comparative cellular analysis of motor cortex in human, marmoset and mouse. Nature 598, 111–119, doi:10.1038/s41586-021-03465-8 (2021).

2 Siletti, K. et al. Transcriptomic diversity of cell types across the adult human brain. Science 382, eadd7046, doi:doi:10.1126/science.add7046 (2023).

3 Johansen, N. et al. Interindividual variation in human cortical cell type abundance and expression. Science 382, eadf2359, doi:doi:10.1126/science.adf2359 (2023).

4 Jorstad, N. L. et al. Comparative transcriptomics reveals human-specific cortical features. Science 382, eade9516, doi:doi:10.1126/science.ade9516 (2023).

5 Werling, D. M. et al. Whole-Genome and RNA Sequencing Reveal Variation and Transcriptomic Coordination in the Developing Human Prefrontal Cortex. Cell Rep 31, 107489, doi:10.1016/j.celrep.2020.03.053 (2020).

6 Mathys, H. et al. Single-cell atlas reveals correlates of high cognitive function, dementia, and resilience to Alzheimer’s disease pathology. Cell 186, 4365–4385. e4327 (2023).

7 Braun, E. et al. Comprehensive cell atlas of the first-trimester developing human brain. Science 382, eadf1226, doi:10.1126/science.adf1226 (2023).

8 Cameron, D. et al. Single-Nuclei RNA Sequencing of 5 Regions of the Human Prenatal Brain Implicates Developing Neuron Populations in Genetic Risk for Schizophrenia. Biol Psychiatry 93, 157–166, doi:10.1016/j.biopsych.2022.06.033 (2023).

9 Hardwick, S. A. et al. Single-nuclei isoform RNA sequencing unlocks barcoded exon connectivity in frozen brain tissue. Nat Biotechnol 40, 1082–1092, doi:10.1038/s41587-022-01231-3 (2022).

10 Herring, C. A. et al. Human prefrontal cortex gene regulatory dynamics from gestation to adulthood at single-cell resolution. Cell 185, 4428–4447.e4428, doi:10.1016/j.cell.2022.09.039 (2022).

11 Morabito, S. et al. Single-nucleus chromatin accessibility and transcriptomic characterization of Alzheimer’s disease. Nat Genet 53, 1143–1155, doi:10.1038/s41588-021-00894-z (2021).

12 Nagy, C. et al. Single-nucleus transcriptomics of the prefrontal cortex in major depressive disorder implicates oligodendrocyte precursor cells and excitatory neurons. Nat Neurosci 23, 771–781, doi:10.1038/s41593-020-0621-y (2020).

13. Biqing, Z. et al. Single-cell transcriptomic and proteomic analysis of Parkinson’s disease Brains. bioRxiv, 2022.2002.2014.480397, doi:10.1101/2022.02.14.480397 (2022).

14 Kang, H. J. et al. Spatio-temporal transcriptome of the human brain. Nature 478, 483–489, doi:10.1038/nature10523 (2011).

15 Wolock, S. L., Lopez, R. & Klein, A. M. Scrublet: Computational Identification of Cell Doublets in Single-Cell Transcriptomic Data. Cell Syst 8, 281–291.e289, doi:10.1016/j.cels.2018.11.005 (2019).

16 Wolf, F. A., Angerer, P. & Theis, F. J. SCANPY: large-scale single-cell gene expression data analysis. Genome Biol 19, 15, doi:10.1186/s13059-017-1382-0 (2018).

17 Lopez, R., Regier, J., Cole, M. B., Jordan, M. I. & Yosef, N. Deep generative modeling for single-cell transcriptomics. Nat Methods 15, 1053–1058, doi:10.1038/s41592-018-0229-2 (2018).

18 Lotfollahi, M. et al. Mapping single-cell data to reference atlases by transfer learning. Nat Biotechnol 40, 121–130, doi:10.1038/s41587-021-01001-7 (2022).

19 Hodge, R. D. et al. Conserved cell types with divergent features in human versus mouse cortex. Nature 573, 61–68, doi:10.1038/s41586-019-1506-7 (2019).

20 Zhu, K. et al. Multi-omic profiling of the developing human cerebral cortex at the single-cell level. Sci Adv 9, eadg3754, doi:10.1126/sciadv.adg3754 (2023).

21 Fu, J. M. et al. Rare coding variation provides insight into the genetic architecture and phenotypic context of autism. Nat Genet 54, 1320–1331, doi:10.1038/s41588-022-01104-0 (2022).

22 Satterstrom, F. K. et al. Large-Scale Exome Sequencing Study Implicates Both Developmental and Functional Changes in the Neurobiology of Autism. Cell 180, 568–584 e523, doi:10.1016/j.cell.2019.12.036 (2020).

23 Meng, X. et al. Multi-ancestry genome-wide association study of major depression aids locus discovery, fine mapping, gene prioritization and causal inference. Nat Genet 56, 222–233, doi:10.1038/s41588-023-01596-4 (2024).

24 Trubetskoy, V. et al. Mapping genomic loci implicates genes and synaptic biology in schizophrenia. Nature 604, 502–508, doi:10.1038/s41586-022-04434-5 (2022).

25 Bellenguez, C. et al. New insights into the genetic etiology of Alzheimer’s disease and related dementias. Nat Genet 54, 412–436, doi:10.1038/s41588-022-01024-z (2022).

26 Bailey, M. H. et al. Comprehensive Characterization of Cancer Driver Genes and Mutations. Cell 173, 371–385.e318, doi:10.1016/j.cell.2018.02.060 (2018).

27 Neftel, C. et al. An Integrative Model of Cellular States, Plasticity, and Genetics for Glioblastoma. Cell 178, 835–849.e821, doi:10.1016/j.cell.2019.06.024 (2019).

28 Wei, X. et al. Integrative analysis of single-cell embryo data reveals transcriptome signatures for the human pre-implantation inner cell mass. Dev Biol 502, 39–49, doi:10.1016/j.ydbio.2023.07.004 (2023).

29 Badia, I. M. P. et al. decoupleR: ensemble of computational methods to infer biological activities from omics data. Bioinform Adv 2, vbac016, doi:10.1093/bioadv/vbac016 (2022).

30 Persad, S. et al. SEACells infers transcriptional and epigenomic cellular states from single-cell genomics data. Nat Biotechnol 41, 1746–1757, doi:10.1038/s41587-023-01716-9 (2023).

31 Van de Sande, B. et al. A scalable SCENIC workflow for single-cell gene regulatory network analysis. Nat Protoc 15, 2247–2276, doi:10.1038/s41596-020-0336-2 (2020).

32 Jassal, B. et al. The reactome pathway knowledgebase. Nucleic Acids Res 48, D498–d503, doi:10.1093/nar/gkz1031 (2020).

33 Escartin, C. et al. Reactive astrocyte nomenclature, definitions, and future directions. Nat Neurosci 24, 312–325, doi:10.1038/s41593-020-00783-4 (2021).

34 Hol, E. M. & Pekny, M. Glial fibrillary acidic protein (GFAP) and the astrocyte intermediate filament system in diseases of the central nervous system. Curr Opin Cell Biol 32, 121–130, doi:10.1016/j.ceb.2015.02.004 (2015).

35 Kang, P. et al. Sox9 and NFIA coordinate a transcriptional regulatory cascade during the initiation of gliogenesis. Neuron 74, 79–94, doi:10.1016/j.neuron.2012.01.024 (2012).

36 Deneen, B. et al. The transcription factor NFIA controls the onset of gliogenesis in the developing spinal cord. Neuron 52, 953–968, doi:10.1016/j.neuron.2006.11.019 (2006).

37 Labreche, K. et al. TCF12 is mutated in anaplastic oligodendroglioma. Nature Communications 6, 7207, doi:10.1038/ncomms8207 (2015).

38 Liu, J., Wang, L. & Li, X. HMGB3 promotes the proliferation and metastasis of glioblastoma and is negatively regulated by miR-200b-3p and miR-200c-3p. Cell Biochem Funct 36, 357–365, doi:10.1002/cbf.3355 (2018).

39 Wang, J. Z. et al. Upregulated YB-1 protein promotes glioblastoma growth through a YB-1/CCT4/mLST8/mTOR pathway. J Clin Invest 132, doi:10.1172/jci146536 (2022).

40 Harris, L. et al. Transcriptional regulation of intermediate progenitor cell generation during hippocampal development. Development 143, 4620–4630, doi:10.1242/dev.140681 (2016).

41 Manojlovic, Z., Earwood, R., Kato, A., Stefanovic, B. & Kato, Y. RFX7 is required for the formation of cilia in the neural tube. Mech Dev 132, 28–37, doi:10.1016/j.mod.2014.02.001 (2014).

42 Tu, S. et al. NitroSynapsin therapy for a mouse MEF2C haploinsufficiency model of human autism. Nature Communications 8, 1488, doi:10.1038/s41467-017-01563-8 (2017).

43 Smith, S. E., Li, J., Garbett, K., Mirnics, K. & Patterson, P. H. Maternal immune activation alters fetal brain development through interleukin-6. J Neurosci 27, 10695–10702, doi:10.1523/jneurosci.2178-07.2007 (2007).

44 Choi, G. B. et al. The maternal interleukin-17a pathway in mice promotes autism-like phenotypes in offspring. Science 351, 933–939, doi:doi:10.1126/science.aad0314 (2016).

45 Shin Yim, Y., et al. Reversing behavioural abnormalities in mice exposed to maternal inflammation. Nature 549, 482–487, doi:10.1038/nature23909 (2017).

46 Kalish, B. T. et al. Maternal immune activation in mice disrupts proteostasis in the fetal brain. Nat Neurosci 24, 204–213, doi:10.1038/s41593-020-00762-9 (2021).

