## Supplementary information for "An integrative single-cell atlas to explore the cellular and temporal specificity of neurological disorder genes during human brain development"

**This file includes:**

Supplementary Figures S1 – S5

Supplementary Tables S1 – S5

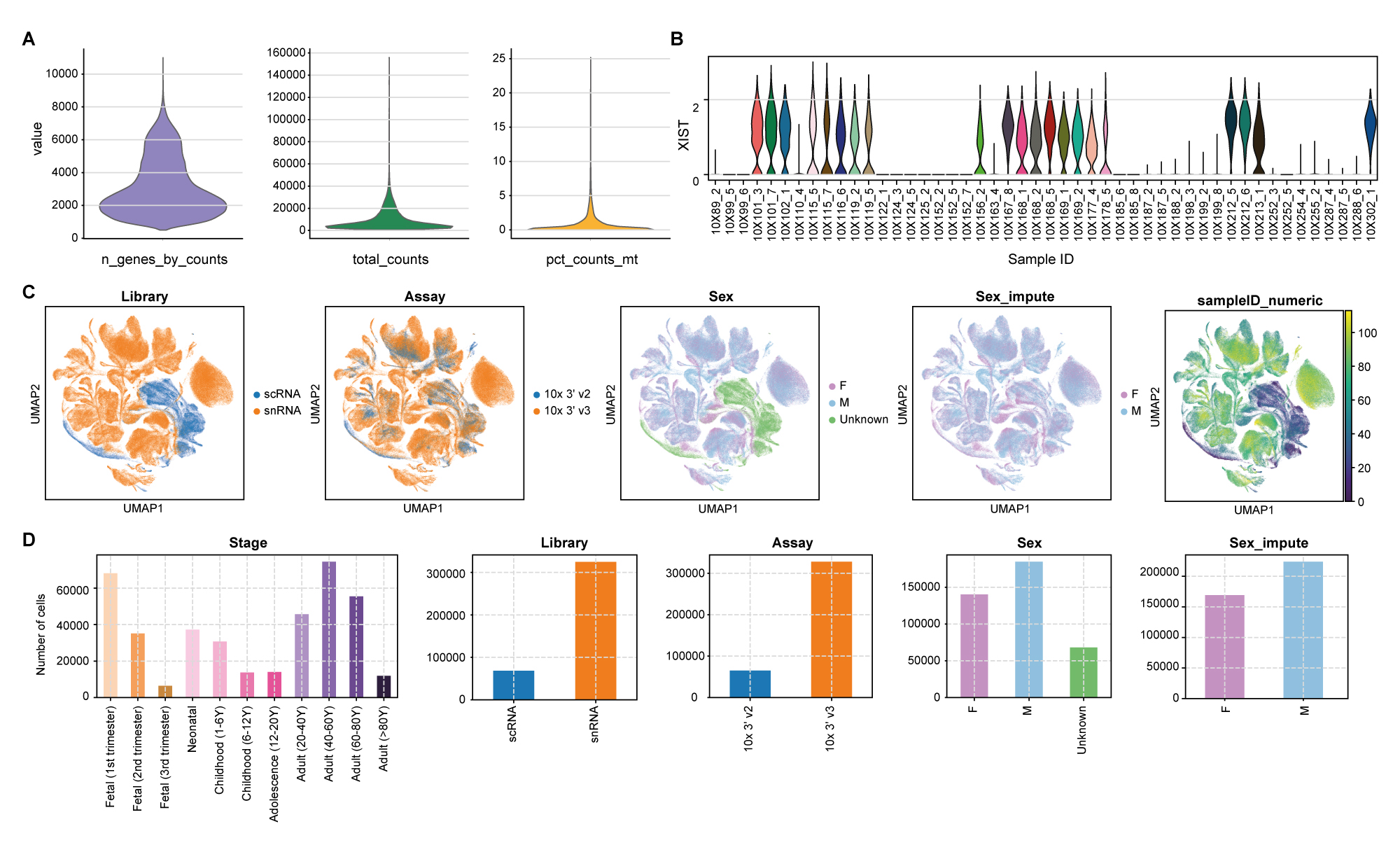

**Supplementary figure 1. Comprehensive overview of quality metrics and sample information in single-cell atlas. A.** Violin plot illustrating quality metrics, including the number of genes with at least 1 count per cell (n_genes_by_counts), the total number of counts per cell (total_counts), and the percentage of counts in mitochondrial genes (pct_counts_mt). **B.** Violin plot for log-normalized XIST expression for samples lacking sex information. **C.** UMAP of the atlas, colored by library, assay, sex, imputed sex (sex_impute), and a numeric representation of sample IDs (sampleID_numeric). **D.** The number of cells categorized by developmental stage, library, assay, sex, and imputed sex.

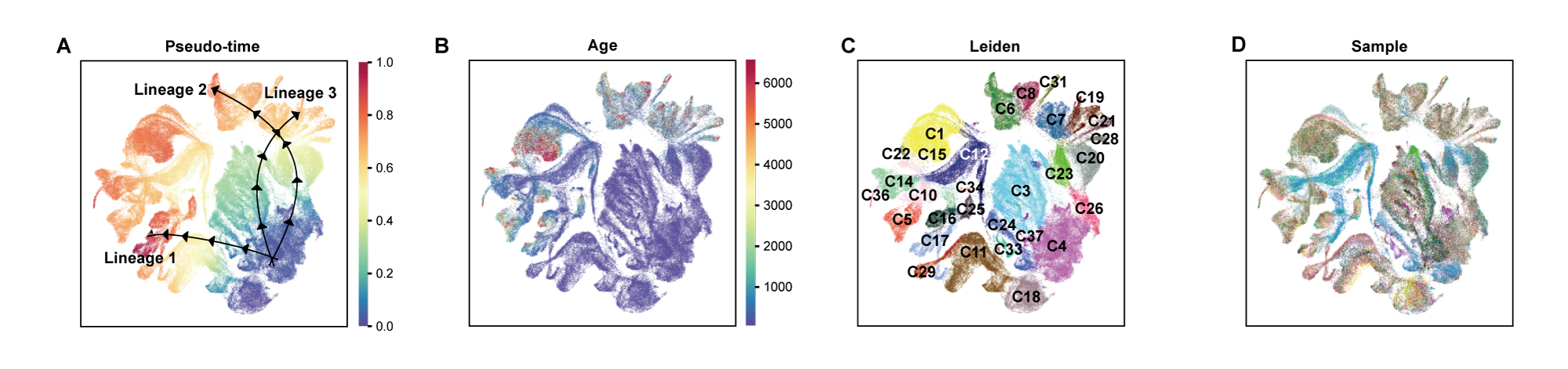

**Supplementary figure 2. Estimated developmental lineages in neuronal cell types.** UMAP visualizations of neuronal cell types colored by (A) estimated pseudotime, (B) gestational age, (C) cluster, and (D) sample ID.

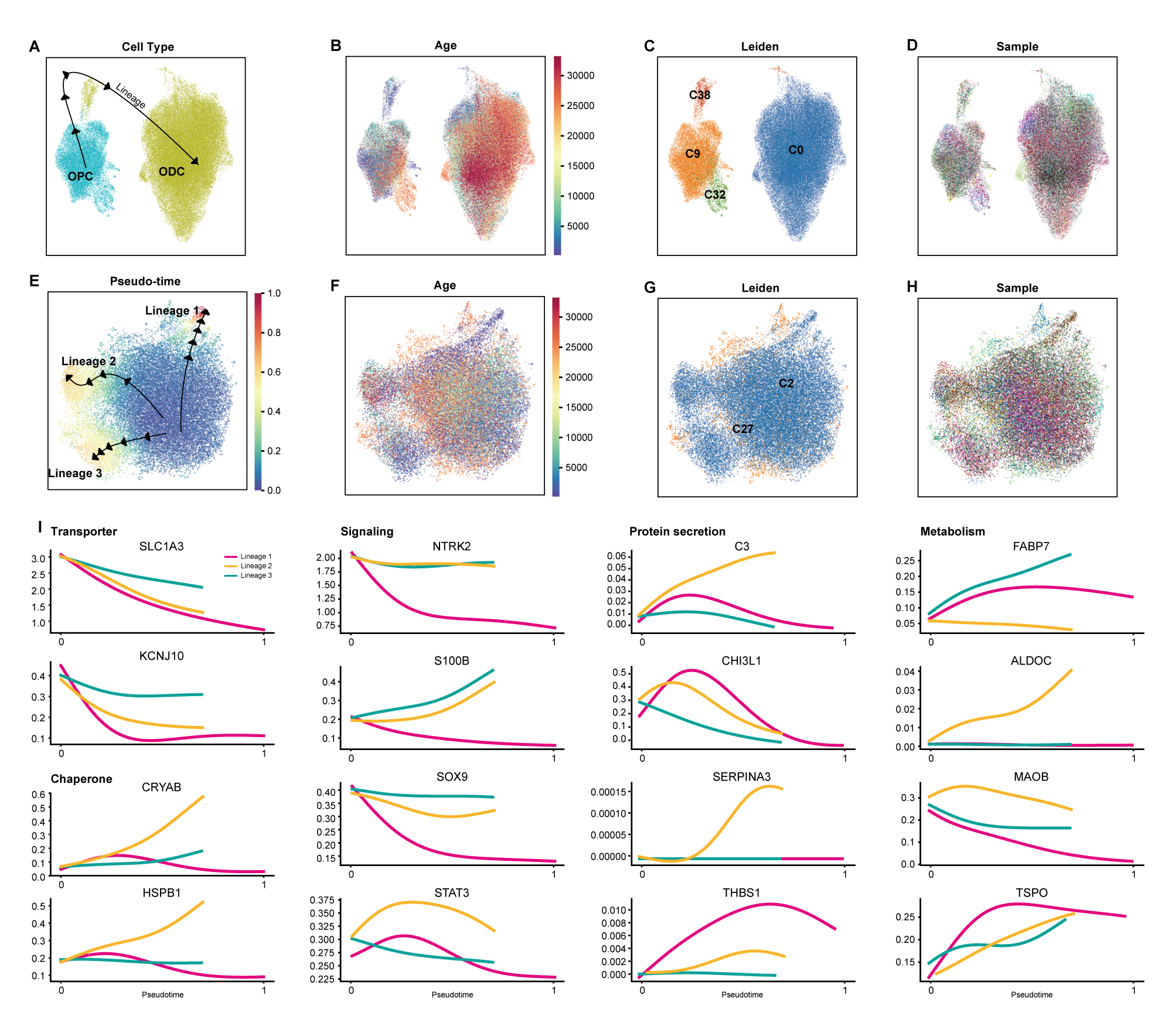

**Supplementary figure 3. Estimated developmental lineages in non-neuronal cell types. A-D.** UMAP visualizations of oligodendrocyte-lineage cell types colored by (A) cell type, (B) gestational age, (C) cluster, and (D) sample ID. **E-H.** UMAP visualizations of astrocytes colored by (A) estimated pseudotime, (B) gestational age, (C) cluster, and (D) sample ID. **I.** Temporal expression patterns of genes associated with function enriched in reactive astrocytes. Expression of genes related to glutamate transporters and K+ channel (Transporter), chaperon activity (Chaperone), transcription factors and receptors, Ca^2+^ binding protein (Signaling), secretion of proteins (Protein secretion), and metabolism including enzyme and lipid transport (Metabolism) of each lineage is dipicted.

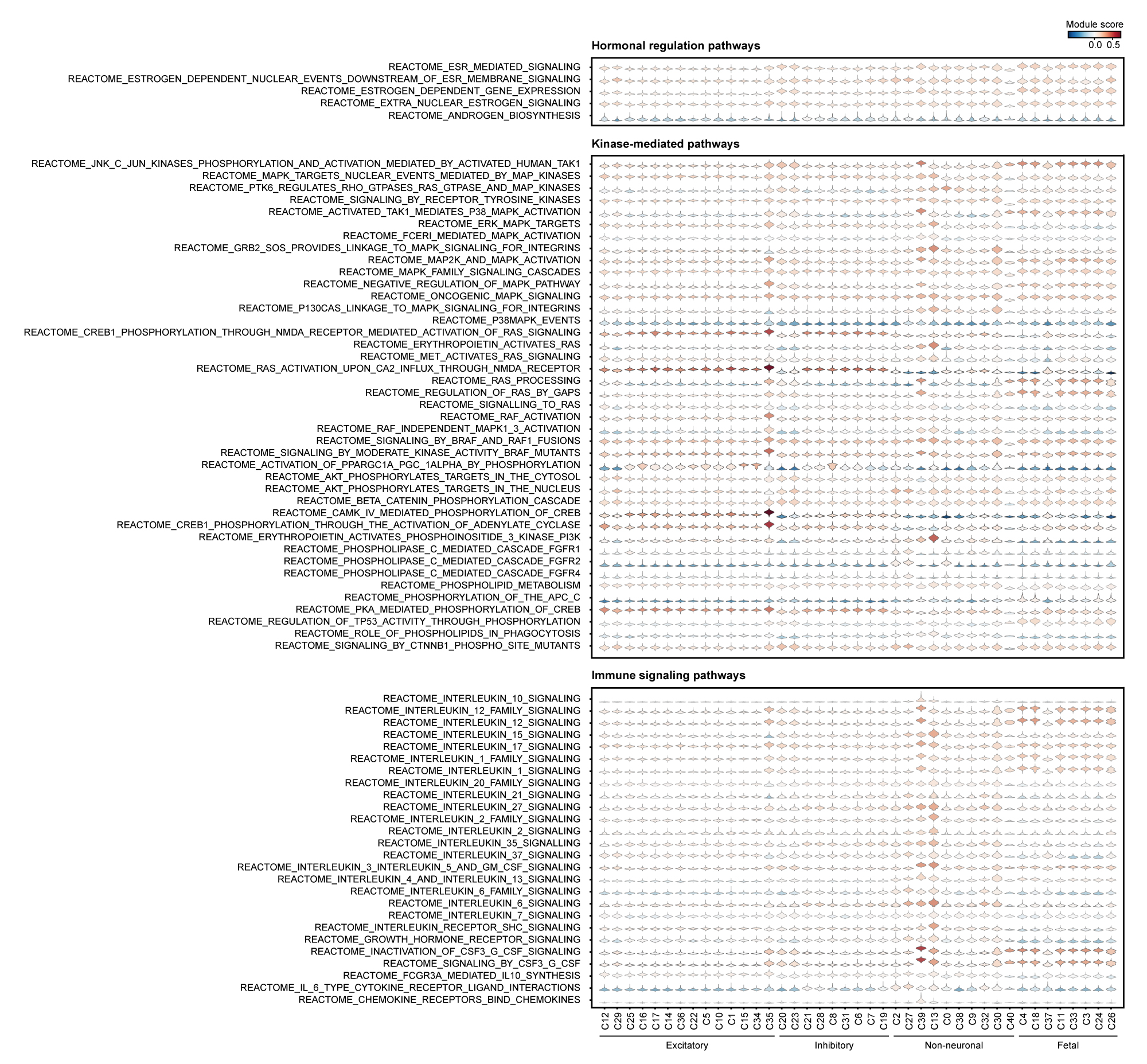

**Supplementary figure 4. Pathway enrichment in early brain development.** Violin plot displaying pathway module scores as the average expression level of pathway genes adjusted for control features.

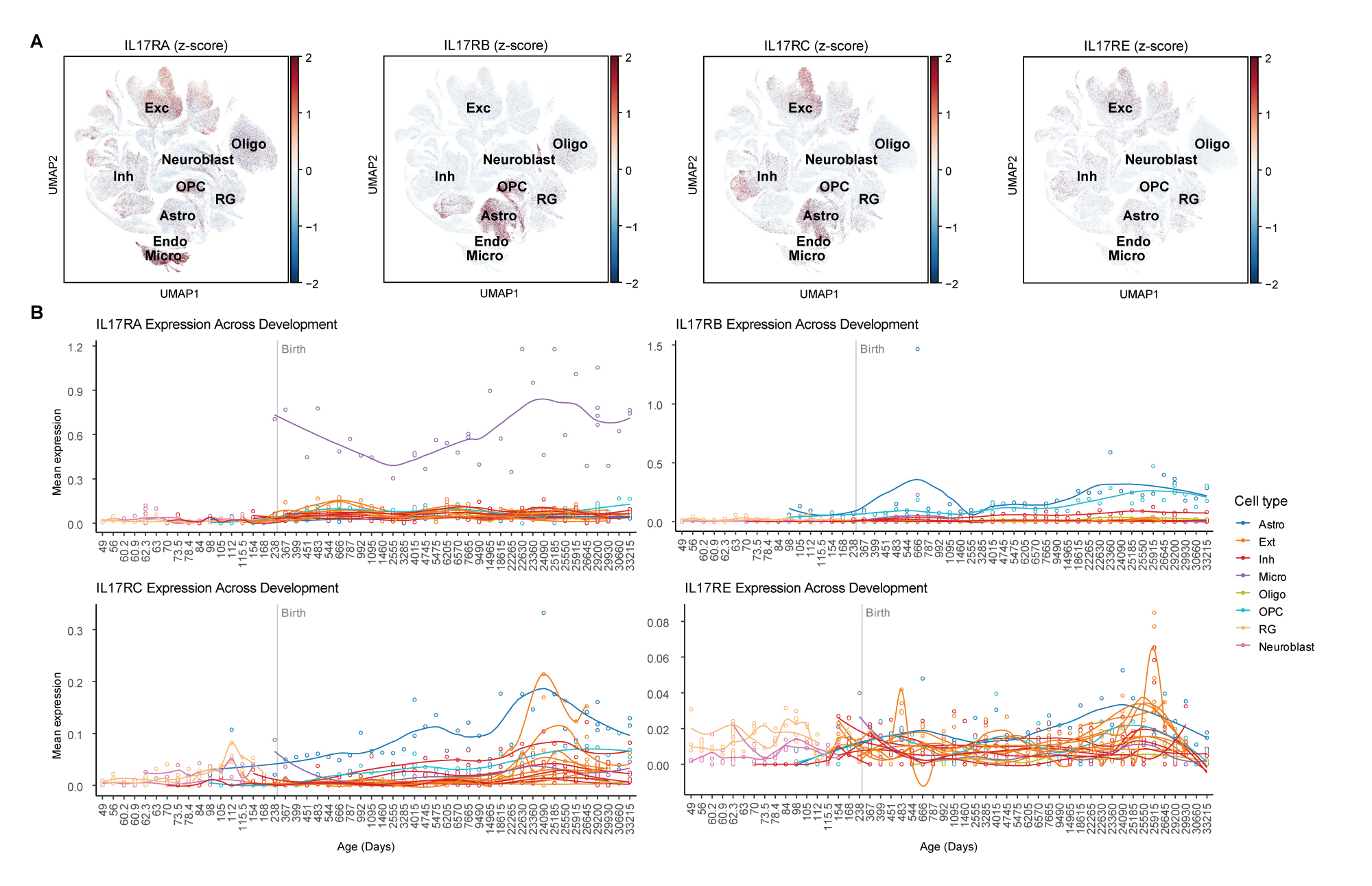
 **Supplementary figure 5. Expression profiles of IL-17 receptor genes (IL17RA, IL17RB, IL17RC, IL17RE). A.** UMAP visualization of z-score normalized IL-17 receptor gene expression. **B.** Expression of IL-17 receptor genes over gestational days. The sample-wise mean of log-normalized gene expression was computed using a pseudo-bulk method. Clusters with at least 4,600 cells (C0-C22) were used.

**Supplementary Tables**

**Supplementary Table S1. Data information**

S1A. Dataset information

S1B. Sample information

**Supplementary Table S2. Identification of cluster-specific DEGs and validation**

S2A. Wilcoxon rank sum test result for cluster-specific DEGs

S2B. Fisher’s exact test with previously identified cell type markers

**Supplementary Table S3. Association of clusters with neurological disorders**

S3A. Neurological disorder genes with details on DEG and lineage associations

S3B. Fisher’s exact test with neurological disorder risk genes

S3C. Fisher’s exact test with cellular states of glioblastoma

**Supplementary Table S4. Pathway enrichment analysis for trajectory lineages**

S4A. Gene Ontology terms enriched for neuronal lineage 1

S4B. Gene Ontology terms enriched for neuronal lineage 2

S4C. Gene Ontology terms enriched for neuronal lineage 3

S4D. Gene Ontology terms enriched for early cells of lineage 1

S4E. Gene Ontology terms enriched for middle cells of lineage 1

S4F. Gene Ontology terms enriched for late cells of lineage 1

S4G. Gene Ontology terms enriched for early cells of oligodendrocyte lineage

S4H. Gene Ontology terms enriched for late cells of oligodendrocyte lineage

S4I. Gene Ontology terms enriched for astrocyte lineage 1

S4J. Gene Ontology terms enriched for astrocyte lineage 2

S4K. Gene Ontology terms enriched for astrocyte lineage 3

**Supplementary Table S5. Multifaceted analysis of regulatory mechanisms**

S5A. Predicted activated transcription factor list per cluster

S5B. Signatures of immune signaling, hormonal regulation, and kinase-mediated pathways

S5C. Wilcoxon rank sum test for cell-type-wise comparison of pathway scores
